# Correlating Synaptic Ultrastructure and Function at the Nanoscale

**DOI:** 10.1101/588848

**Authors:** Lydia Maus, Bekir Altas, JeongSeop Rhee, Nils Brose, Cordelia Imig, Benjamin H. Cooper

**Affiliations:** Department of Molecular Neurobiology, Max Planck Institute of Experimental Medicine, 37075 Göttingen, Germany; University of Göttingen, 37073 Göttingen, Germany; Department of Pharmacology, University of Maryland School of Medicine, Baltimore, MD 21201

## Abstract

Despite similarities in the composition of the molecular release machinery, synapses can exhibit strikingly different functional transmitter release properties and short- and long-term plasticity characteristics. To address the question whether ultrastructural differences could contribute to this functional synaptic heterogeneity, we employed a combination of hippocampal organotypic slice cultures, high-pressure freezing, freeze substitution, and 3D-electron tomography to resolve the spatial organization of vesicle pools at individual active zones (AZ) in two functionally distinct synapses, namely Schaffer collateral (SC) and mossy fiber (MF) synapses. We found that mature MF and SC synapses harbor equal numbers of docked vesicles at their AZs, MF synapses at rest exhibit a second pool of possibly ‘tethered’ vesicles in the AZ vicinity, and MF synapses contain at least three morphological types of docked vesicles, indicating that differences in the ultrastructural organization of MF and SC synapses may contribute to their respective functional properties and corresponding plasticity characteristics.

## INTRODUCTION

Transmitter release at presynaptic active zones (AZs) is triggered by membrane depolarization, typically in the form of an action potential (AP), and the concomitant influx of Ca^2+^ via voltage-gated Ca^2+^ channels (VGCCs). The Ca^2+^ signal is then detected by sensor proteins, which elicit the Soluble NSF Attachment Protein Receptor (SNARE)-mediated fusion of docked and primed, fusion-competent synaptic vesicles (SVs) (Südhof, 2013). Although the various types of synapses in the mammalian brain employ strikingly similar sets of proteins for transmitter release, their functional characteristics can differ dramatically, particularly the initial probability of SV fusion (P_r_) and its short-term plasticity (STP) during and after trains of multiple APs (Regehr, 2012). In extreme cases, certain synapse types, particularly those with high initial P_r_, have a ‘phasic’ character and exhibit strong synaptic depression during AP trains, while others, often ones with low initial P_r_, are ‘tonic’ in nature and show strong frequency facilitation (Neher and Brose, 2018). These presynaptic features, in turn, are critical determinants of synaptic computing and play key roles in many brain processes (Regehr, 2012).

P_r_ and STP are affected by multiple factors, including the AP shape, the type, density, and modulation of presynaptic VGCCs, the distance between VGCCs and primed SVs, the type and concentration of presynaptic Ca^2+^-buffers, the type of exocytotic Ca^2+^-sensor, the types of SNARE-proteins involved in SV fusion, or the clearance of SV material from AZ fusion sites (Dittman and Ryan, 2009; Kaeser and Regehr, 2017). Importantly in this context, the size of the readily-releasable pool (RRP) of primed SVs at rest and the rate of RRP exhaustion and refilling during ongoing stimulation are parameters that synapses employ and adapt purposively to shape STP (Neher and Brose, 2018; Regehr, 2012).

EM-tomography analyses of high-pressure frozen, cryo-substituted hippocampal Schaffer collateral (SC) synapses showed that primed SVs are in point contact with the AZ membrane and that this state depends on Munc13s and all three SNARE proteins (Imig et al., 2014; Siksou et al., 2009). Complementary functional evidence indicates that this docked and primed state of SVs is reversible and can be generated within only a few milliseconds (Chang et al., 2018; He et al., 2017; Miki et al., 2018). This led to the notion of a ‘loosely docked and primed SV state’ (LS) that that can be rapidly converted into a ‘tightly docked and primed SV state’ (TS), in which SVs can fuse readily upon the arrival of an AP. In terms of synapse function, the notion of LS-SVs and TS-SVs and their rapid interconversion was proposed to explain key features of phasic, depressing and tonic, facilitating synapses, where phasic synapses are characterized by a large TS-SV pool at rest, which is exhausted with ongoing stimulation to cause synaptic depression, while tonic synapses feature an initially small TS-SV pool that is progressively filled in an activity- and Ca^2+^-dependent manner during ongoing stimulation to cause frequency facilitation (Neher and Brose, 2018). This model of LS-SV vs. TS-SVs has similarities with functional definitions of heterogeneous SV pools, e.g. ‘reluctantly/slowly’ vs. ‘fully/rapidly’ releasable (Lee et al., 2012; Neher, 2015; Neher and Brose, 2018) or ‘primed’ vs. ‘superprimed’ SVs (Taschenberger et al., 2016) as well as with morphological classifications of membrane-proximal SV pools such as tethered vs. docked SVs (Imig et al., 2014; Watanabe et al., 2013).

The present study was performed to test the prediction that defined functional features of different presynapse types have an ultrastructural basis, and — more concretely — that the phasic vs. tonic nature of synapses might become manifest as a difference in the relative proportion of tethered and docked SVs. We used EM-tomography of high-pressure frozen, cryo-substituted hippocampal organotypic slices to study low-P_r_, (Lawrence et al., 2004) strongly facilitating (Salin et al., 1996) mossy fiber (MF) synapses, which had not been studied at a level of resolution required for an accurate discrimination of morphologically and functionally distinct SV pools, and to compare them to SC synapses, which have a heterogeneous, but on average substantially higher P_r_ (Helassa et al., 2018; Oertner et al., 2002). Our data indicate that differences in the spatial organization of AZ-proximal SVs, rather than the absolute number of docked and primed SVs at AZs, contributes to the STP differences between SC and MF synapses.

## RESULTS

### Intact MF Pathway in Hippocampal Organotypic Slice Cultures

To test whether SC and MF synapses exhibit distinct ultrastructural features that could explain their respective functional properties and STP characteristics, we combined hippocampal organotypic slice cultures, high-pressure freezing (HPF), automated freeze substitution (AFS), and electron-tomography (ET) on plastic sections to study the distribution of SVs at individual AZs with nanometer precision (Imig et al., 2014; Studer et al., 2014). For the comparative ultrastructural analysis of distinct hippocampal synapse types, organotypic slices offer multiple advantages: (i) Local hippocampal circuits (Figure S1) and synaptic properties, including short-term plasticity characteristics, remain largely intact (Galimberti et al., 2006; Mori et al., 2004; De Simoni et al., 2003), (ii) the functional and ultrastructural properties of mature synaptic connections that are difficult to reinstate in dissociated cultures can be analyzed in an *in-situ*-like environment, even with perinatally lethal mouse mutants (Imig et al., 2014), (iii) cultured slices recover from trauma induced during sectioning, and (iv) slice dimensions are compatible with HPF cryofixation, yielding a near-native preservation of synaptic ultrastructure (Figures 1 and S1) (Imig and Cooper, 2017).

**Figure 1.**
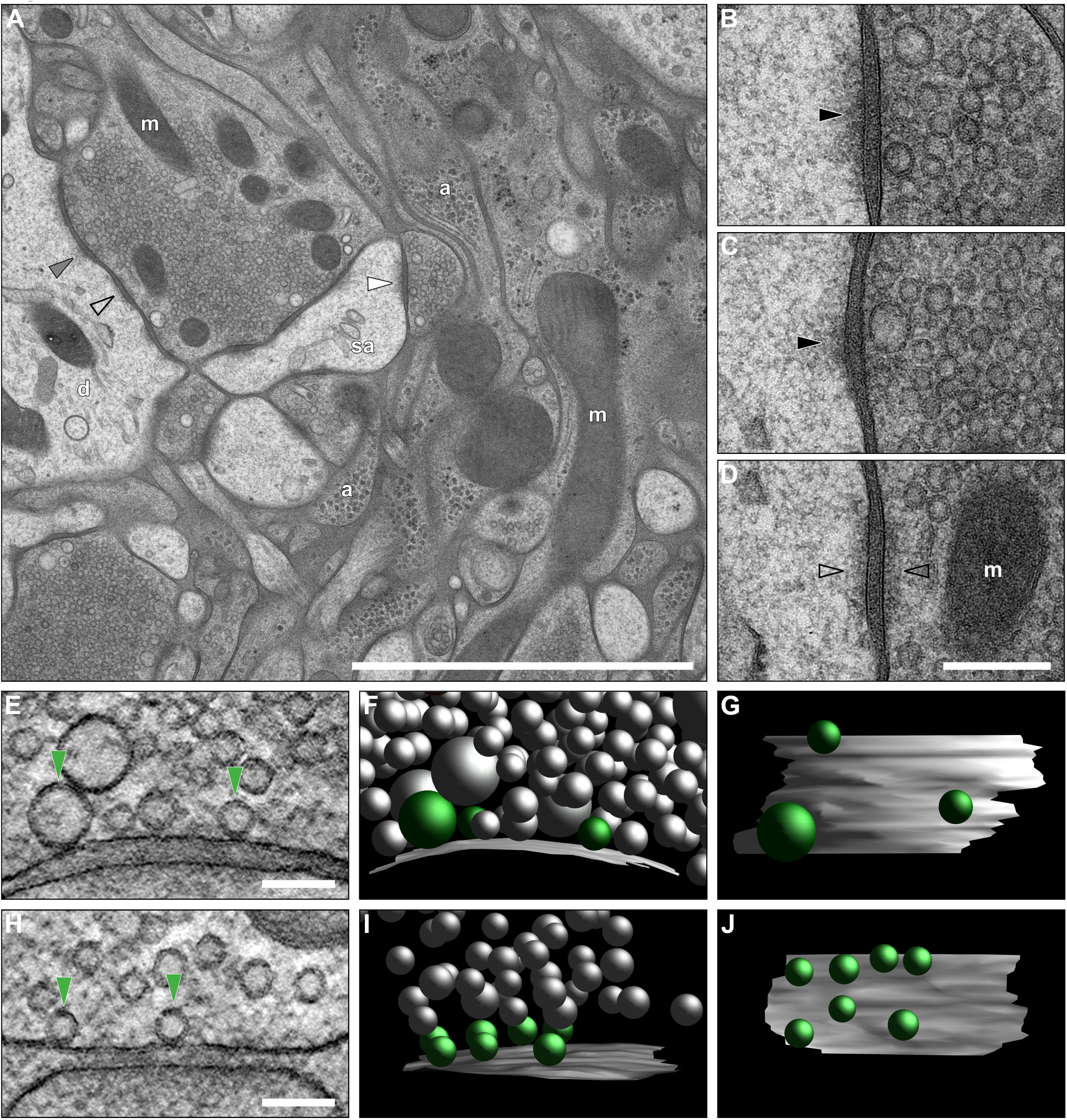
Ultrastructural Characterization of the MF-CA3 PC Synapse in Organotypic Hippocampal Slice Cultures Prepared by HPF and AFS. (**A**) 2D-Electron micrograph of a MFB forming multiple synaptic contacts with a postsynaptic CA3 PC. MF synapse characteristics include a large presynaptic bouton densely packed with synaptic vesicles (SVs) and multiple postsynaptic densities (PSDs). (**B**-**D**, enlarged from **A**) MFBs form three types of contacts with CA3 PCs: Asymmetric spine (**B**, white arrowhead in **A**) and dendritic synapses (**C**, grey arrowhead in **A**) (black arrowheads, PSDs), as well as *puncta adherentia* onto dendritic shafts (**D**, black arrowhead in **A**), which are characterized by symmetrical paramembranous electron-dense material (open arrowheads). (**E**, **H**) ET subvolumes of MF (**E**) and SC (**H**) synapses (docked vesicles indicated by green arrowheads). (**F**, **I**) 3D models of synaptic profiles from MF (**F**) and SC (**I**) synapses (AZ, grey; docked vesicles, green; nonattached vesicles, gray). (**G**, **J**) Orthogonal views of MF (**G**) and SC (**J**) AZs. Abbreviations: a, astrocytic processes; d, dendrite; m, mitochondria; sa, spine apparatus Scale bars: A, 2 µm; B-D, 200 nm; E-J, 100 nm.

Light microscopic examination of cultured slices revealed Synaptoporin-positive MF boutons (MFBs) (Singec et al., 2002) in the *stratum lucidum* clustered near primary dendrites of MAP-2-positive CA3 pyramidal neurons (Figure S1B) and equipped with multiple AZ release sites as indicated by the high density of colocalized Bassoon puncta (Figure S1D). Biocytin-filled CA3 pyramidal neurons displayed complex, multi-compartmental spines (thorny excrescences; Figure S1D), the morphological hallmark of hippocampal MF synapses (Chicurel and Harris, 1992). Electron micrographs from ultrathin sections (Figures 1A, S1E, and S1G) showed large MFBs with excellent ultrastructural preservation and a gross morphology comparable to that of perfusion-fixed (Figures S1F, S1I, and S1J) and acute slice preparations (Figure S1H). MFB-CA3 pyramidal cell (PC) synaptic contacts are established on complex spines (Figure 1B) and dendritic shafts (Figure 1C). *Puncta adherentia* (Figure 1D) between MFBs and CA3 PCs were observed exclusively in contact with dendritic shafts and were difficult to differentiate from AZs with a low SV-occupancy in thick sections prepared for ET. We therefore restricted our comparative ET analysis of SC and MF AZ organization to release sites onto dendritic spines.

ET analysis of synaptic sub-volumes from SC (Supplementary Video 1) and MF (Supplementary Video 2) AZs allowed us to precisely measure distances between SVs and the AZ membrane (Figures 1E-1J). For the analysis of MF-CA3 PC AZs, ET is especially helpful as spines often exhibit complex shapes and highly convoluted membranes, which may confound the accurate detection of SV docking in 2D projection images acquired from ultrathin sections (Imig and Cooper, 2017). SVs with no measurable distance between the SV and plasma membrane lipid bilayers (0-2 nm) were considered ‘docked’ (Figures 1E and 1H) (Imig and Cooper, 2017; Imig et al., 2014).

### Comparison of SC and MF Synapses in Hippocampal Slice Cultures

We selected two developmental time points for our study, days *in vitro* (DIV)14 and DIV28, to cover a period of significant morphological (Amaral and Dent, 1981) and functional maturation, such as an increase in excitatory postsynaptic current (EPSC) amplitudes, an enhanced degree of low frequency facilitation at 1 Hz in mice (Marchal and Mulle, 2004), and a shift from long-term depression to potentiation after high-frequency stimulation (Battistin and Cherubini, 1994), and to correlate our findings with existing ultrastructural (Chicurel and Harris, 1992; Rollenhagen et al., 2007; Studer et al., 2014) and functional (Hallermann et al., 2003; Jonas et al., 1993; Midorikawa and Sakaba, 2017) datasets on MF synapses. At DIV14 and DIV28, we observed three morphological vesicle types docked at MF AZ membranes (Figure 2C, E, H-J): (i) small clear-core SVs (33-55 nm Ø; Figure 2A), (ii) ‘giant’ clear-core vesicles (GVs, 60-120 nm Ø; Figure 2B), and (iii) dense-core vesicles (DCVs, 46-91 nm Ø; Figure 2C). Neither GVs nor DCVs docked at SC AZs in any condition analyzed.

**Figure 2.**
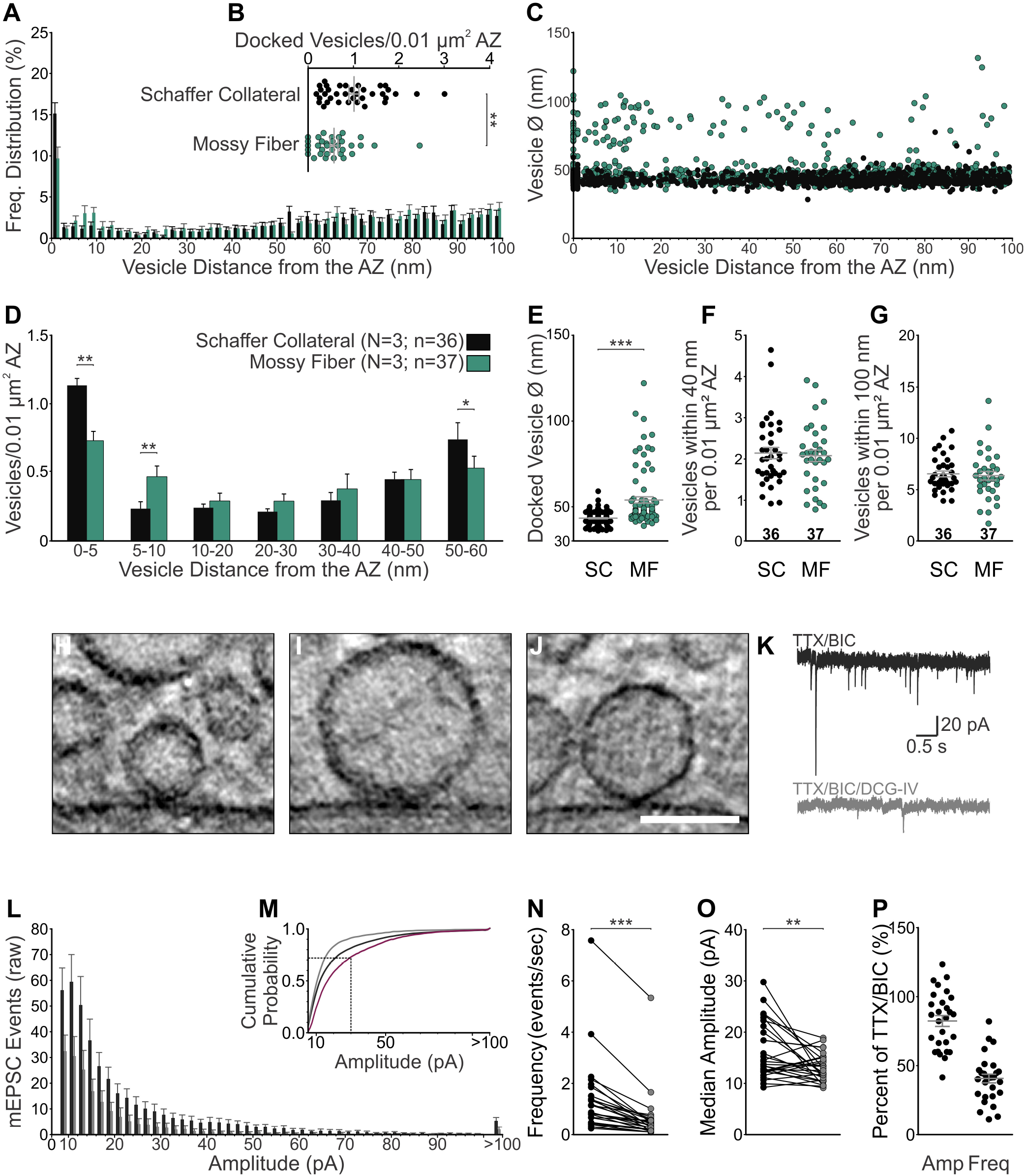
Morphological and Functional Dissection of Distinct Vesicle Pools at MF-CA3 PC Synapses in Organotypic Hippocampal Slice Cultures at DIV14. (**A**-**G**) Morphological Characterization of MF and SC AZs (N = 3 cultures, SC n = 36, and MF n = 33 tomograms). See Table S1A (**A**) Spatial distribution of vesicles within 100 nm of the active zone (AZ) membrane in SC and MF synapses. (**B**) Scatterplot of the mean number of docked, clear-cored vesicles (SVs and GVs) normalized to AZ area. (**C**) Plot of vesicle diameters for all vesicles analyzed and their respective distance to the AZ membrane. (**D**) Mean number of vesicles within bins of 5 nm and 10 nm from the AZ normalized to AZ area. (**E**) Scatterplot of SV diameters for all docked vesicles analyzed in SC (n = 116 vesicles) and MF (n = 81) synapses. (**F**, **G**) Scatterplots of vesicles within 40 nm (**F**) and 100 nm (**G**) of the AZ membrane normalized to AZ area. (**H**-**J**) Virtual slices of three morphologically distinct vesicle types docked at MF synapses: SVs (**H**), GVs (diameter > 60 nm; **I**), and DCVs (**J**). (**K**-**P**) Effects of DCG-IV on mEPSCs recorded from CA3 PCs in slice cultures (N = 2 cultures, n = 28 cells). See Table S1G (**K**) Representative traces of mEPSC events recorded from CA3 PCs in the presence of 1 µM tetrodotoxin (TTX) and 10 µM Bicuculline (BIC) to block GABA_A_-receptor mediated events throughout the recording before and after the application of 2 µM DCG-IV. (**L**) Histogram for mEPSC event amplitudes (≥ 8 pA; 2 pA bins and last bin all events ≥ 100 pA) before (black, TTX/BIC) and after application of DCG-IV (grey, TTX/BIC/DCG-IV). Bars represent number of mEPSC events recorded for an indicated amplitude range in a 5 min interval (mean events per cell + SEM). (**M**) Cumulative probability plot of mEPSC amplitudes. The subtracted cumulative frequency distribution of mEPSCs removed by DCG-IV application is indicated in purple and the dotted line marks an mEPSC amplitude of 30 pA (see methods). (**N**) Mean frequency of mEPSCs before and after application of DCG-IV. (**O**) Median amplitudes of mEPSCs before and after application of DCG-IV. (**P**) Scatterplot for relative changes in mean amplitude and frequency after the application of DCG-IV normalized to the control condition (TTX/BIC only). Scale bars: H-J, 100 nm. Values indicate mean ± SEM; *p<0.05; **p<0.01; ***p<0.001.

At DIV14, the number of clear-core vesicles docked to MF AZs normalized to the AZ area was 43% lower than for SC synapses in the same slice (Figures 2B and 2D; Table S1A) although the numbers of vesicles within 40 nm (membrane-proximal) and 100 nm of the AZ were comparable (Figures 2F and 2G; Table S1A). However, at DIV28 (Figures 3 and S2; Table S1B and S1D) the density of docked vesicles at MF AZs was comparable to that in SC synapses at DIV14 (Figure 2; Table S1A) and DIV28 (Imig et al., 2014). This developmental increase in the density of docked vesicles in MFBs accompanies previously reported morphological changes during MF maturation between P14 and 21, including the generation of additional and more complex postsynaptic spines, the emergence of spine apparatus, an enlargement of the presynaptic terminal, and an increase in synaptic release site number and size (Amaral and Dent, 1981). Thus, our findings contribute to the understanding of the functional changes seen during MF maturation in this time period.

**Figure 3.**
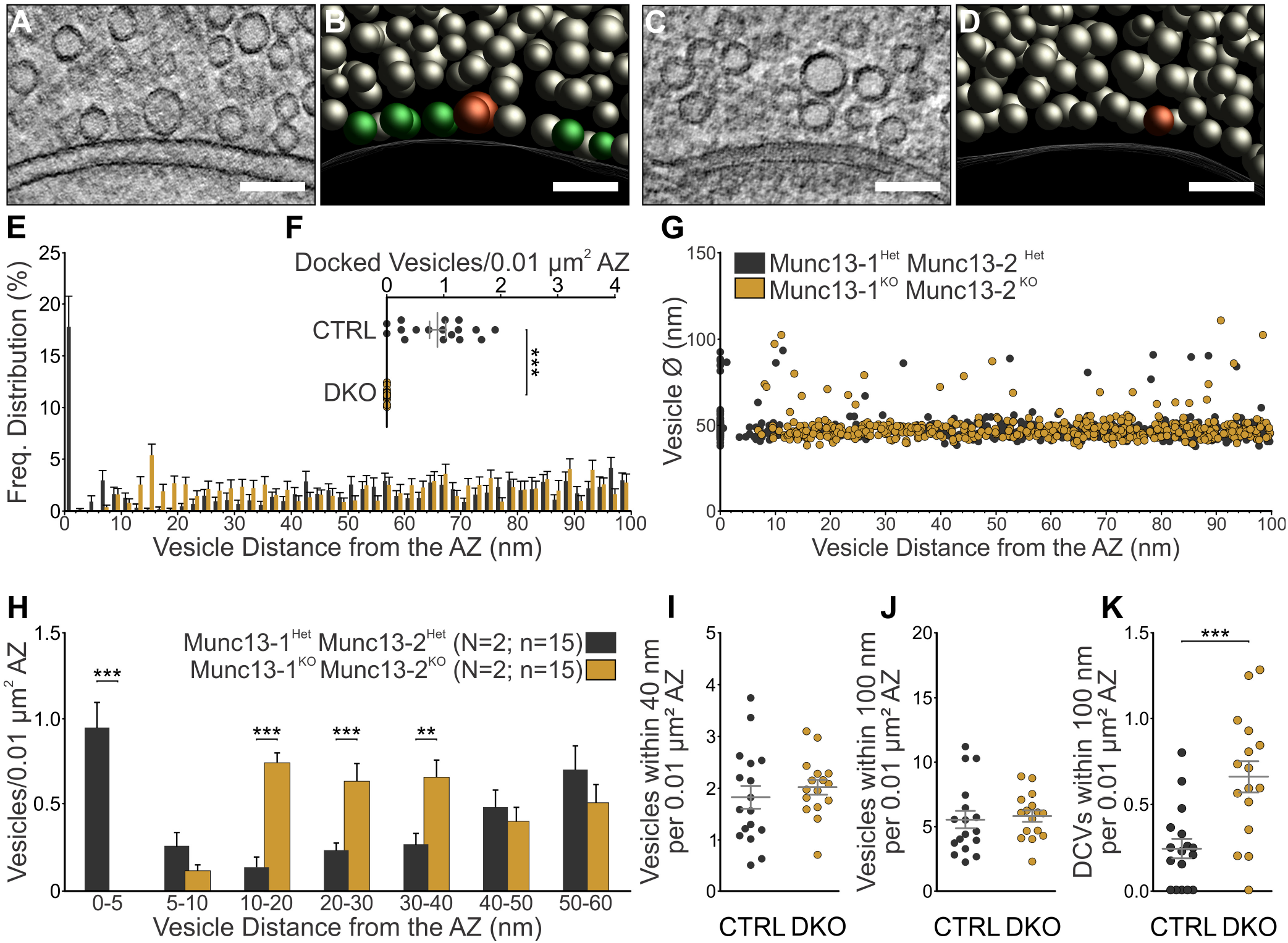
Electron Tomographic Analysis of Vesicle Pools in M13 DKO and CTRL MF Synapses. (**A**, **C**) ET subvolumes of CTRL (**A**) and M13 DKO (**C**) MF synapses. (**B**, **D**) 3D models of CTRL (**B**) and M13 DKO (**D**) MF synapses. (**E**) Spatial distribution of vesicles within 100 nm of the AZ membrane (N = 2 cultures; n = 15 tomograms). See Table S1B (**F**) Scatterplot of docked vesicles in M13 CTRL and DKO MF synapses normalized to AZ area. (**G**) Plot of vesicle diameters for all vesicles analyzed and their respective distance to the AZ membrane. (**H**) Mean number of vesicles within bins of 5 nm and 10 nm from the AZ normalized to AZ area. (**I**, **J**) Scatterplots of vesicles within 40 nm (**G**) and 100 nm (**H**) of the AZ membrane normalized to AZ area. (**K**) Scatterplot of DCVs within 100 nm of the AZ membrane normalized to AZ area. Scale bars: A-D, 100 nm. Values indicate mean ± SEM; *p<0.05; **p<0.01; ***p<0.001.

Interestingly, and unlike SC synapses, MF synapses at both developmental time points exhibited a second pool of membrane-proximal SVs accumulated at ∼5-15 nm from the AZ (Figures 2A, 2D, S2C, and S2F), reminiscent of the tethered SV pool in docking- and priming-deficient mutant SC synapses (Imig et al., 2014; Siksou et al., 2009). These are adequately positioned to undergo rapid priming to sustain the RRP of docked SVs during repetitive stimulation, which is in accord with the notion that tonic synapses at rest harbor an abundant supply of tethered or LS-SVs (Neher and Brose, 2018). However, the similar densities of docked SVs in mature (DIV28) SC (Imig et al., 2014) and MF synapses (Figure S2D) indicate that the differences between these synapse types as regards initial P_r_ and STP are not dictated by the availability of docked and primed or TS-SVs (Neher and Brose, 2018).

### GVs in MFBs Are Not a Mere By-Product of Synaptic Activity

The ultrastructural profile of MF AZs in cryo-fixed organotypic slices were compared with those of (i) acutely dissected and cryo-fixed hippocampal slices (Figures S2I-S2O) and of (ii) mice transcardially perfused according to two different protocols used previously in seminal EM studies on rat MFBs (Chicurel and Harris, 1992; Rollenhagen et al., 2007) (Figures S2A-S2H; Table S1D). GVs were observed in all preparations, albeit in slightly lower numbers in aldehyde-fixed (Figure S2E) as compared to cryo-fixed material (Figures 2C, 2E, S2K, and S2M; Table S1D), indicating that their presence is not a slice-culture-specific phenomenon. Strikingly, aldehyde-fixed MFBs exhibited a general depletion of docked and tethered SVs and GVs (Figures S2A-S2H), that correlated with an increased abundance of Ω-shaped fusion intermediates (Figures S2A and S2B), and a redistribution of AZ-proximal vesicles (Figures S2C and S2F) comparable to previous observations in mouse somatosensory cortex (Korogod et al., 2015). Although the extent of these effects depends on fixative composition and osmolarity (Figures S2D and S2F-S2H; Table S1D), our data indicate that aldehyde fixation perturbs presynaptic ultrastructure and can thus confound the functional interpretation of morphological data.

GVs in MFBs may represent precursor vesicles of somatic origin, endocytic intermediates of local SV recycling, or neurotransmitter-filled SV-type end-products of a specialized mode of presynaptic SV biogenesis. To distinguish between these possibilities, we investigated the activity-dependence of GV abundance. To this end, we analyzed MF AZs from Munc13-1/2-deficient (M13 DKO) mice, in which hippocampal synaptic transmission (Varoqueaux et al., 2002) and exocytosis-coupled ultrafast endocytosis near AZs (Watanabe et al., 2013) are completely abolished (Figure 3). In M13-DKO MFBs, we observed a complete loss of SV, GV, and DCV docking (Figures 3A-3F) but no changes in the number of vesicles within 40 or 100 nm from the AZ (Figures 3G and 3H; Table S1B), which is in accord with previous data on SC synapses (Imig et al., 2014; Siksou et al., 2009). Unexpectedly, we observed a three-fold increase in the number of DCVs in the AZ vicinity (Figure 3I; Supplementary Video 3; Table S1B). Importantly, GVs were still observed in the vicinity of M13-DKO MF AZs (Figures 3C, D and 3G). Similarly, acute pharmacological blockade of slice culture network activity (TTX/NBQX/AP-V) or presynaptic MF activity (TTX/DCG-IV) failed to eliminate GVs from MFB AZs (Figure S3; Table S1F). These data demonstrate that GVs are especially abundant in MF-CA3 synapses, that their generation is largely independent of synaptic activity and SV recycling, and that Munc13s are absolutely required for SV, GV, and DCV docking at MF AZs.

### GVs as the Morphological Correlates of Giant mEPSCs

It was suggested previously that GVs in MF synapses contain glutamate and represent the morphological correlates of large-amplitude miniature (m)EPSCs (giant mEPSCs; ≥100 pA) recorded from CA3 PCs (Henze et al., 2002). We measured spontaneous mEPSCs in CA3 PCs in slice culture preparation and confirmed the presence of such large mEPSCs of ≥100 pA (Figure 2L). Upon selective blockade of MF glutamate release by the mGluR2 agonist DCG-IV (Kamiya et al., 1996), we observed a strong reduction in the mean mEPSC frequency (Figures 2N and 2P) as well as a reduced frequency of large-amplitude mEPSCs. The latter effect was reflected by reductions in median and mean mEPSC amplitudes (Figures 2O and 2P) and by a change in the cumulative mEPSC amplitude distribution (Figure 2M). These data are in line with the fact that lesioning neonatal rat dentate gyrus granule cells by γ-irradiation to deplete MF inputs to CA3 PCs causes a loss of large-amplitude mEPSCs in CA3 PCs (Henze et al., 1997).

To correlate the amplitude distribution of recorded mEPSCs with the observed size distribution of docked SVs, we assumed that (i) a 10 pA mEPSC (the mode of the DCG IV-sensitive mEPSC amplitude distribution) reflects the quantal glutamate release from a SV with an outer Ø of 44 nm (the mode of the docked vesicle size distribution), (ii) the intravesicular glutamate concentration is independent of vesicle volume, and (iii) that postsynaptic glutamate receptor saturation at MF-CA3 synapses is negligible during a single mEPSC. We determined that the fusion of an SV with an outer Ø of 60 nm (size threshold for classification as GV) would generate an mEPSC of ∼30 pA, and a GV with an outer Ø of 85 nm (mean Ø of all docked vesicles with Ø>60 nm) would correspond to an mEPSC of ∼98 pA (see methods). The cumulative frequency distribution of DCG-IV-sensitive mEPSC amplitudes (Figure 2M, purple line) indicates that approximately 27% of all mEPSCs are ≥30 pA and could therefore originate from glutamate quanta released from GVs (Ø>60 nm), which is in good agreement with our finding that GVs comprise 20% of clear-core vesicles docked at MF AZs (Figure 2E). These findings provide further support for the notion that GVs at MF AZs are indeed the morphological correlate of the large mEPSC events observed at MF synapses. Beyond this, it cannot be ruled out that some giant mEPSCs are caused by spontaneous multiquantal release from non-MF inputs onto the same PC, which would explain the fact that not all mEPSCs larger than 100 pA are blocked by DCG-IV.

### cAMP-Dependent Changes in Transmitter Release Do Not Affect SV Docking

In MF synapses, changes in cAMP levels alter presynaptic strength (Kamiya and Yamamoto, 1997) and trigger a presynaptic form of long-term synaptic plasticity via downstream effectors such as PKA (Tzounopoulos et al., 1998; Weisskopf et al., 1994). To test if corresponding cAMP signaling reorganizes AZ-proximal SV pools, we applied the adenylyl cyclase activator forskolin (FSK) or the presynaptic mGluR2 agonist DCG-IV to organotypic slices prior to HPF to potentiate or depress neurotransmitter release from MF synapses. Neither FSK nor DCG-IV affected docked or AZ-proximal SV and GV pools at MF synapses (Figures 4A-4G; Table S1C). This is in agreement with the finding that increased cAMP levels in dissociated MF terminals only had a moderate effect on the RRP size (Midorikawa and Sakaba, 2017). Interestingly though, the number of docked DCVs in FSK-treated MF synapses was two-fold and six-fold increased compared to vehicle control and DCG-IV-treated slices, respectively (Table S1C). We detected only a small and non-significant tendency towards an increased SV docking in FSK treated slices (Figures 4A, 4B, and 4E; Table S1C), so that a greater proportion of SVs within 0-40 nm from the AZ were attached to the AZ membrane (Figures 4F and 4G). In principle, this is in line with the notion that increased cAMP/PKA signaling increases P_r_ and the efficacy of SV fusion via improved coupling to VGCCs (Midorikawa and Sakaba, 2017). However, given the rather subtle morphological changes we observed, we favor an interpretation where cAMP-mediated PKA activation affects proteins of the SV fusion machinery at a post-docking/priming step to facilitate SV fusion, e.g. via phosphorylation of Complexins (Cho et al., 2015). Such a scenario would also explain the enhanced apparent Ca^2+^-sensitivity of SV fusion after FSK application (Midorikawa and Sakaba, 2017).

**Figure 4.**
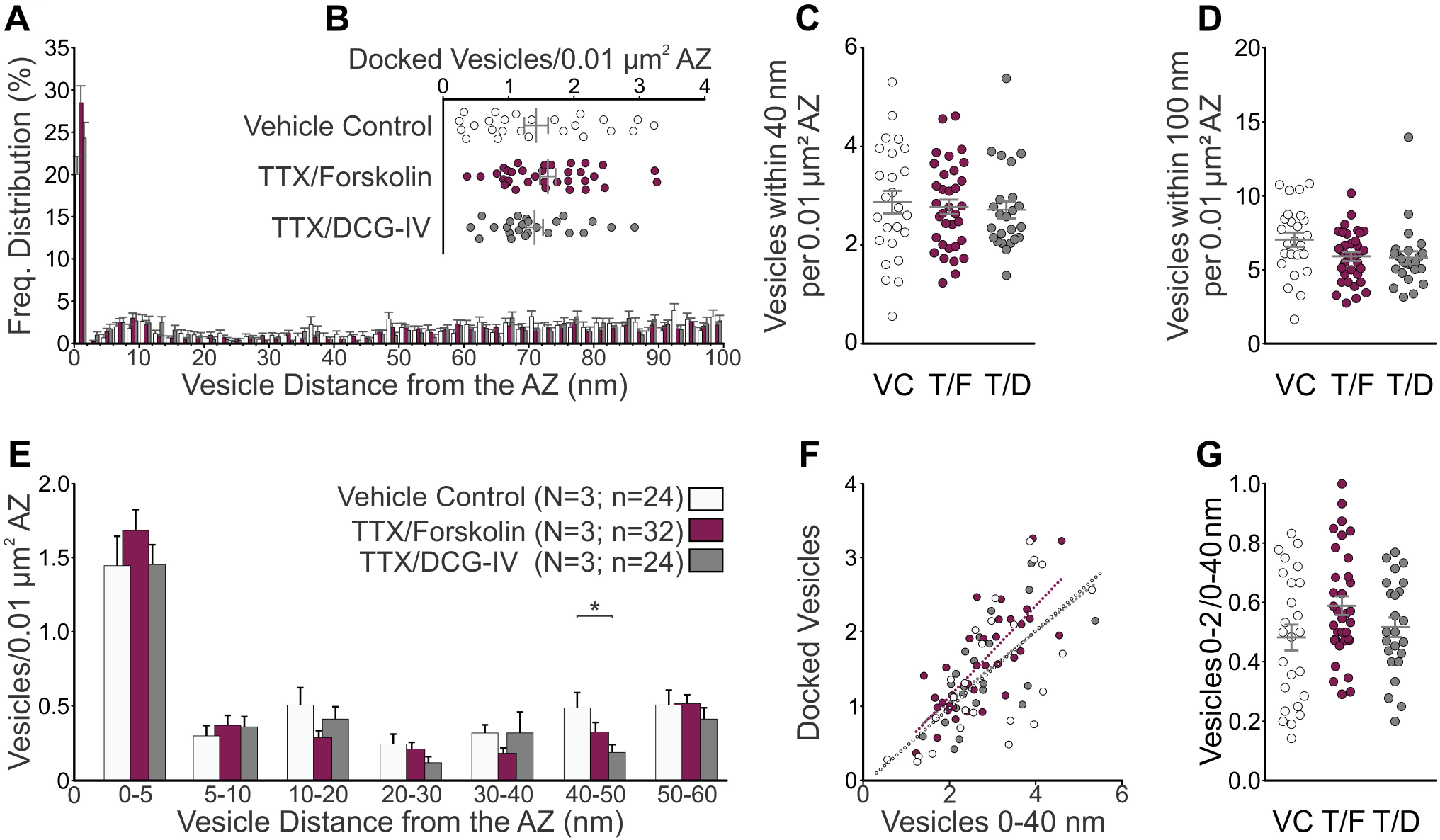
Pharmacological Manipulation of Presynaptic cAMP Levels. (**A**) Spatial distribution of vesicles within 100 nm of the AZ membrane in MF synapses treated for 15 min with either vehicle control (VC; 1 µM TTX; N = 3 cultures; n = 24 tomograms), TTX and 2 µM DCG-IV (T/D; N = 3; n = 24) or TTX and 25 µM forskolin (T/F; N = 3; n = 32). See Table S1C (**B**) Scatterplot of docked vesicles normalized to AZ area. (**C**, **D**) Scatterplots of vesicles within 40 nm (**C**) and 100 nm (**D**) normalized to AZ area. (**E**) Mean number of vesicles within bins of 5 and 10 nm from the AZ normalized to AZ area. (**F**) Number of docked vesicles normalized to 0.01 µm^2^ AZ area plotted as a function of the number of vesicles within 40 nm of the AZ normalized to 0.01 µm^2^ AZ area (VC, y=0.517x-0.0612 (R^2^=0.434); T/D, y=0.470x+0.123 (R^2^=0.423); and T/F, y=0.610x-0.091 (R^2^=0.600); linear regression test for difference). (**G**) Ratio of docked vesicles (0-2 nm) to vesicles within 40 nm of AZ membrane. Values indicate mean ± SEM; *p<0.05; **p<0.01; ***p<0.001.

## DISCUSSION

### Methodology

We combined HPF, AFS, and ET to investigate whether the spatial organization of SVs at presynaptic AZs shapes the distinct transmitter release and STP properties of hippocampal SC and MF synapses. In the course of our study, we made the important methodological observation that standard chemical fixatives deplete docked SV pools, probably at least partly due to the increased osmolarity of aldehyde-supplemented buffers (Figures S2A-S2I), although we cannot exclude the possibility that more refined or optimized fixation conditions may cause less severe effects on presynaptic vesicle pools. This finding, which matches previous morphological (Korogod et al., 2015) and functional observations (Smith and Reese, 1980), does not merely highlight a fundamental risk involved in comparing EM results obtained based on different experimental parameters (e.g. fixation methods, imaging approaches, and/or docking criteria). Rather, it indicates (i) that EM data obtained with chemically fixed samples, while very useful in numerous contexts, are not suitable for the functional interpretation of morphologically defined presynaptic SV pools, and (ii) that a combination of HPF, AFS, and ET, as employed in the present study, or cryo-EM, provided that hyperosmolarity problems can be circumvented, should be used instead.

### Synapse Ultrastructure and STP

The mechanistic basis of the strong frequency facilitation of MFBs, a hallmark of this synapse (Salin et al., 1996) that co-determines its role as a ‘conditional detonator’ synapse (Vyleta et al., 2016), has long been enigmatic (Nicoll and Schmitz, 2005). Recent evidence indicates that multiple factors converge to shape P_r_ and STP at MF synapses. They include a large coupling distance (∼70 nm) between VGCCs and SVs (Vyleta and Jonas, 2014), action potential broadening (Geiger and Jonas, 2000), Ca^2+^ buffer saturation (Blatow et al., 2003; Vyleta and Jonas, 2014), and specialized exocytotic Ca^2+^-sensors (Jackman et al., 2016).

A key aim of the present study was to determine whether differences in the nanometer-scale distribution of SV pools at AZs also contribute to the distinct transmitter release and STP properties of MF terminals, as compared to the higher-P_r_ SC synapses in the same circuit. We made three key observations in this context: (i) Mature MF and SC synapses have equal numbers of docked SVs at their AZs (Figure S2; Table S2A and S2D) and (Imig et al., 2014), indicating that the low initial P_r_ in MF synapses is not caused by a limited availability of primed and docked SVs at release sites. (ii) In contrast to SC synapses, MF synapses at rest exhibit a second pool of membrane-proximal, possibly ‘tethered’ SVs in the AZ vicinity (Figures 2A and 2D), which are well positioned to be rapidly mobilized in a Ca^2+^-dependent manner during sustained synaptic activity and contribute to the prominent facilitation at MF synapses. (iii) MFBs contain at least three morphological vesicle types (SVs, GVs, DCVs) that all dock to presynaptic AZs in a Munc13-dependent manner (Figure 3), indicating that a proper molecular composition of release sites and the core neuronal priming machinery are essential prerequisites for vesicle fusion in MFBs.

### P_r_, RRP, and Docked SVs

The issue as to whether the number of docked SVs per AZ can predict the P_r_ of a given synapse and its STP characteristics is the focus of substantial interest and controversy (Éltes et al., 2017; Neher and Brose, 2018; Xu-Friedman and Regehr, 2004). At its core is the question as to whether docked SVs comprise the RRP of functionally primed, fusion-competent SVs (Kaeser and Regehr, 2017; Neher and Brose, 2018; Xu-Friedman and Regehr, 2004). The biggest current conundrum in this context is the fact that the RRP is a relatively vague concept, usually defined functionally, in terms of SVs that can fuse in response to a given stimulus in a given synapse, and most frequently measured using postsynaptic responses as a proxy (Kaeser and Regehr, 2017; Neher, 2015; Neher and Brose, 2018).

We calculated mean docked vesicle numbers per AZ and per MFB (∼320 SVs, 37 GVs, 15 DCVs) as well as the mean membrane surface area for each vesicle type in mature (DIV28) MF synapses (see methods). Assuming a specific membrane capacitance of 1 μF/cm^2^ (Hallermann et al., 2003), fusion of all docked vesicles, irrespective of their type, would change the membrane capacitance (ΔC_m_) by ∼32 fF per MFB. By including the second pool of AZ-proximal, tethered vesicles into the analysis to test the notion that vesicles in an LS or tethered state can rapidly be converted in a Ca^2+^-facilitated manner into a TS, docked, and fully-primed state (Chang et al., 2018; Neher and Brose, 2018), the fusion of all vesicles within 0-40 nm of the AZ would correspond to a ΔC_m_ of ∼88 fF per MFB. Previous studies showed that long depolarizations (30-100 ms) evoke a ΔC_m_ of 50-100 fF in entire MFBs, which corresponds, depending on the SV Ø used for conversion, to a functional RRP of 500-1400 SVs per MFB or ∼40 SVs per AZ (Hallermann et al., 2003; Midorikawa and Sakaba, 2017). This matches our calculations for the combined pools of docked and AZ-tethered vesicles (ΔC_m_ of ∼88 fF) and earlier estimates of the number of SVs with centers within 60 nm of the AZ (Rollenhagen et al., 2007). That functionally obtained RRP estimates at MFBs are much larger than the number of docked vesicles we found, confirms the problem that some ‘pool-depleting’ stimulation protocols used to assess the number of primed SVs at a given synapse are not sufficiently refined to dissect the relative contributions of docked vs. tethered SV pools and cannot account for fast priming during stimulation (Neher, 2015; Neher and Brose, 2018). In essence, long depolarisations of low-P_r_ MF synapses may not only evoke fusion of docked vesicles, but additionally induce fast and Ca^2+^-mediated priming of tethered vesicles and their subsequent fusion.

### SVs, GVs, and DCVs at MF AZs

Our study revealed the presence of docked SVs, GVs, and DCVs at AZs as a unique feature of MFBs (Figure 2H-J), and showed a loss of docking of all three vesicle types upon deletion of Munc13 family vesicle priming proteins (Figure 3D; Table S1B). These data indicate that Munc13s are not only required for SV docking and priming (Imig et al., 2014; Siksou et al., 2009), but also for the docking and priming of GVs and DCVs at MF AZs. As regards to MF GVs, which we never detected in any other synapse type, but which have previously been reported in aldehyde-perfused MFBs (Henze et al., 2002; Laatsch and Cowan, 1966; Rollenhagen et al., 2007), our data (Figure 2) lead to the conclusion that GVs at MF synapses represent neurotransmitter-containing vesicles that can fuse and release their content. However, their origin remains unknown and additional or even alternative functional contexts need to be considered. Although some synapse types exhibit compound SV fusion during stimulation, generating giant vesicular structures and corresponding increases in mEPSC amplitudes (He et al., 2009), the persistence of GVs in genetically and pharmacologically silenced presynapses in organotypic slices (Figures 3 and S3) argues against compound fusion or compensatory endocytic membrane-retrieval (Watanabe et al., 2013) as the primary mode of GV formation in MFBs. However, we cannot rule out the possibility that endocytic processes contribute to a subpopulation of GVs in MFBs. Alternatively, GVs might originate from DCVs that have undergone non-collapse fusion, degranulation, and rapid retrieval (Laatsch and Cowan, 1966), a phenomenon observed during neuroendocrine DCV fusion (Shin et al., 2018). Indeed, filamentous electron dense material is occasionally observed in the lumen of MF GVs (e.g. Figure 1E). However, the prominent accumulation of DCVs in Munc13-deficient samples implies that DCV fusion in MFBs is dramatically impaired, yet GVs with and without filamentous luminal content remain, indicating that GV formation does not critically depend on synaptic membrane, SV, GV, or DCV cycling activity. A final possibility is that at least some of the GVs in MFBs represent vesicles of somatic origin that are trafficked via anterograde axonal transport from dentate granule cells. Consistent with this notion, tomographic reconstructions of granule cell axons in the *stratum lucidum* of acute hippocampal slices revealed a range of trafficked vesicle types, including large, clear-cored vesicles (Figure S3). However, it is unclear whether and how such unusual precursor vesicles would be appropriately equipped to participate in synaptic signaling. In any case, MF GVs are highly unlikely to be of artifactual nature and do not appear to be a mere peculiarity without function, so that further cell biological and functional studies into their role at MF synapses seem worthwhile.

## ACKNOWLEDGMENTS

This work was supported by the German Research Foundation (SFB1286/A01, B.H.C.), and the European Commission (ERC Advanced Grant SynPrime; N.B.). We are grateful to Peter Jonas, Noemi Holderith, and Vincent O’Connor for helpful comments on the manuscript, Holger Taschenberger for discussions and advice, and Manuela Schwark for excellent technical support.

## AUTHOR CONTRIBUTIONS

Conceptualization, N.B., C.I., B.H.C.; Methodology, C.I., B.H.C.; Formal Analysis, L.M.; Investigation, L.M., B.A., C.I., B.H.C.; Resources, J.R., N.B.; Writing – Original Draft, L.M., N.B., C.I., B.H.C.; Writing – Review and Editing, all authors; Visualization, L.M., C.I., B.H.C.; Supervision, N.B., C.I., B.H.C.; Funding Acquisition, N.B., B.H.C.

## MATERIALS AND METHODS

### Experimental Model and Subject Details

#### Mouse Breeding

Mouse breeding and transcardial perfusion was done with permission of the Niedersachsisches Landesamt für Verbraucherschutz und Lebensmittelsicherheit (LAVES; 33.19.42502-04-15/1817 and 33.19-42502-04-18/2756). All animals were kept according to the European Union Directive 63/2010/EU and ETS 123. All wild-type animals (WT) used in this study were C57BL/6N. Mice were housed in individually ventilated cages (type II superlong, 435 cm^2^ floor area; TECHNIPLAST) under a 12 h light/dark cycle at 21 ± 1°C with food and water *ad libitum*. The health status of the animals was checked regularly by animal care technicians and a veterinarian. Mice lacking Munc13-1 (Unc13A) and Munc13-2 (Unc13B) (Augustin et al., 1999; Varoqueaux et al., 2002) and control (CTRL) littermates were generated from crossing animals with the genotype Unc13A^+/−^ (Munc13-1) Unc13B^+/−^ (Munc13-2) with Unc13A^+/−^ Unc13B^−/−^. CTRL animals with the genotypes Unc13A^+/−^ Unc13B^+/−^ and Unc13A^+/+^ Unc13B^+/−^ were used for tomographic analysis.

#### Tissue Culture

Hippocampal organotypic slice cultures were prepared using the interface method (Stoppini et al., 1991) according to previously published protocols (Imig et al., 2014; Studer et al., 2014). Slices were prepared from WT animals postnatal day (P) 3-7 and from M13 DKO and littermate CTRL animals at embryonic day 18 (E18) due to the severe perinatally lethal phenotype (Varoqueaux et al., 2002). Pregnant females at gestational stage E18 were anaesthetized and decapitated and embryos removed by hysterectomy. Pups were decapitated and the brain was quickly removed and placed into preparation medium (97 mL Hank’s balanced salt solution, 2.5 ml 20% glucose, and 1 ml 100 mM kynurenic acid, pH adjusted to 7.4). Both hippocampi were dissected and transferred with the entorhinal cortex attached onto a tissue chopper platform. Three hundred µm-thick hippocampal slices were cut perpendicular to the longitudinal axis of the hippocampus using a McIlwain tissue chopper and then quickly washed off the stage into preparation medium. Slices were carefully transferred onto sterile Millipore membrane confetti pieces that were placed on top of 6-well membrane inserts in culture medium (22.44 mL ddH2O, 25 mL 2xMEM, 25 mL BME, 1 mL GlutaMAX, 1.56 mL 40% Glucose, 25 mL horse serum). Residual preparation medium was removed from the inserts using a P200 pipette. A maximum of 4 hippocampal slices were cultured per membrane insert. Slice culture medium was changed 24 hours after preparation and then 2-3 times per week for the remaining culture period. Slices were cultured for either 14 or 28 days at 37°C and 5% CO_2_. In line with previous observations, we did not observe neurogenesis in the dentate gyrus in organotypic slices cultured in the presence of serum (Raineteau et al., 2004), as assessed by the lack of calretinin-positive immature cells (Brandt et al., 2003) in the sub-granular zone of the dentate gyrus (Figure S3).

#### Method Details

##### Experimental Design

Experiments from WT mice were performed on 3-4 independent slice cultures and from M13 DKO and CTRL cultures on two independent cultures due to the severity of the phenotype. The following time points were included into the analysis: DIV14 (refers to culture experiments performed on DIV13-16), DIV28 (DIV27/28), P18 (refers to acute slice preparations performed on P17/P18), and P28 (refers to transcardial perfusions performed on P27/P28). The P18 time point for the acute preparations was chosen to match the exact age of WT DIV14 slice cultures that were prepared on P3-P6 (see below). Each perfusion protocol was performed on two WT animals. Animals of both genders were analyzed. The experimenter was blinded for the experiments that involved pharmacological treatments of organotypic slices. Electrophysiological recordings from CA3 PCs and morphological analyses from MFBs were performed in the CA3b,c regions of the hippocampus.

##### HPF, AFS, and EM Sample Preparation

###### Transcardial Perfusion

WT animals at P28 were given a lethal dose of Avertin (2,2,2,-Tribromoethanol) via intraperitoneal injection. Deeply anaesthetized animals were transcardially perfused first with 0.9% sodium chloride solution and then one of two fixatives [*Fixative 1 (approximately 1900 mOsm* (Hayat, 1981)): Ice-cold 4% paraformaldehyde (PFA), 2.5% glutaraldehyde (GA) in 0.1 M phosphate buffer (PB), pH 7.4 (Rollenhagen et al., 2007); *Fixative 2 (approximately 1200 mOsm* (Hayat, 1981)): 37°C 2% PFA, 2.5% GA, 2 mM CaCl_2_, in 0.1 M cacodylate buffer (CB) (Chicurel and Harris, 1992)].Fixative osmolalities were determined from literature (Hayat, 2000). Brains were removed from the animals and post-fixed in respective fixative overnight at 4°C. The brains were washed in ice-cold 0.1 M PB pH 7.4. Hundred-µm coronal hippocampal sections were cut using a Leica Vibratome (Leica VT 1200S, amplitude of 1.5 mm, cutting speed 0.1 mm/sec) in 0.1 M PB pH 7.4. Sections were stored in ice-cold 0.1 M PB and high-pressure frozen on the same day.

###### Acute brain slice preparation

Anaesthetized WT animals aged P18 were quickly decapitated and the brains were removed from the skull. One hippocampus was quickly dissected and placed onto a McIlwain tissue chopper stage. Two hundred-µm thick slices were cut perpendicular to the longitudinal axis of the hippocampus. Slices were washed into a petri dish containing 20% BSA in HEPES-buffered artificial cerebrospinal fluid (ACSF). Slices were separated and the CA3 and CA1 regions were punched out of the hippocampal slice using a 1.5 mm diameter biopsy punch. The hippocampal regions were loaded into 3 mm aluminium planchettes (Leica Cat# 1677141 for type A and 1677142 for type B) and the remaining space was filled with cryoprotectant (20% BSA in ACSF) and immediately cryofixed within 5 min of decapitation. We did not attempt to recover slices after sectioning, because previous studies have demonstrated that prolonged incubation in ACSF deteriorates the freezing quality of brain tissue (Korogod et al., 2015; response to the reviewers). Only synapses that lacked morphological evidence of extensive stimulation (i.e. depletion of SVs in the terminal, endocytic pits) were analyzed.

###### HPF of organotypic slices

*Slices* were changed to fresh slice culture medium 24 hours before HPF. Immediately prior to freezing, untreated slices were transferred into warm, pre-equilibrated slice culture medium and excess membrane confetti were trimmed away from each slice. Slices were then transferred into non-penetrating 20% BSA cryoprotectant dissolved in culture medium. Slices were loaded into aluminium specimen carriers (type A, Leica Cat# 16770126, outer diameter 6 mm, inner cavity depth 100 µm) membrane-side up and filled with cryoprotectant. The filled carriers were loaded into middle plates of the HPF sample holder. The flat side of type-B aluminium carriers (Leica Cat# 16770127) were coated with 1-hexadecene and placed flat-side down onto the sample-filled carrier to serve as “lids”. Excess liquid was removed with Whatman filter paper. The sample holder was then quickly assembled and loaded into the HPF device (Leica HPM 100). Cryo-fixed samples were stored in liquid nitrogen until further processing.

###### Pharmacological silencing experiments

The protocol for acute pharmacological silencing of cultured slices was based on a previously published protocol for the application of drugs to organotypic slice cultures (Studer et al., 2014). Briefly, DIV14 organotypic slices were placed onto a new, sterile membrane inserts in a 6-well plate containing fresh organotypic slice culture medium as well as different receptor and channel blockers: i) Vehicle control (VC, organotypic slice culture medium only); ii) T/N/A (medium containing 1 µM tetrodotoxin (TTX), 2 µM 2,3-Dioxo-6-nitro-1,2,3,4-tetrahydrobenzo[*f*]quinoxaline-7-sulfonamide (NBQX), and 50 µM D-(-)-2-Amino-5-phosphonopentanoic acid (D-AP5)); iii) T/D (medium containing 1 µM TTX and 2 µM (2*S*,2’*R*,3’*R*)-2-(2’,3’-Dicarboxycyclopropyl)glycine (DCG-IV)). Then, 50 µL drops containing T/N/A, T/D, or VC medium were pipetted onto slices and incubated at 37°C and 5% CO_2_ before slices were prepared for HPF. The medium and cryoprotectant for subsequent steps (see above) always contained the respective drugs. Slices were frozen 10 min after drug exposure.

###### Pharmacological manipulation of presynaptic cAMP levels

DIV28 organotypic slices were transferred onto a new membrane insert in a six-well plate containing fresh organotypic slice culture medium: i) Vehicle control (1 µM TTX, ddH_2_O, and DMSO); ii) T/D (1 µM TTX, 2 µM DCG-IV, and DMSO); iii) T/F (1 µM TTX, ddH_2_O, and 25 µM forskolin). Slices were placed back in the incubator, exposed to the pharmacological treatment for 15 min, and were then again taken out of the incubator and prepared for HPF (see above).

###### AFS

Automated freeze substitution (AFS) was performed as previously described (Imig et al., 2014; Rostaing et al., 2006). Briefly, samples were incubated in 0.1% tannic acid in anhydrous acetone for 4 days at −90°C and then fixed with 2% osmium tetroxide in anhydrous acetone with the temperature slowly ramping up to 4°C over several days. Samples were washed in acetone and brought to room temperature for EPON infiltration and embedding. Ultimately, gel-capsules filled with 100% EPON were inverted on the sample carriers and polymerized at 60°C for 24-48 h. Polymerized blocks were removed from the glass slides and the aluminium carriers were carefully trimmed off of the blocks. With the exception of culture slices from E18 mice that fit into 3 mm aluminium carriers, blocks were further trimmed down to fit an EM grid.

###### Ultramicrotomy

Five hundred nm-thick sections were cut on a Leica Ultracut UCT ultramicrotome until tissue appeared and cell body lamination became apparent in semithick sections. Then, 4 to 5 200 nm-thick semithin sections were cut and collected onto Formvar-filmed, carbon-coated and glow-discharged copper mesh grids for electron tomography. Subsequently, a few ultrathin sections (60 nm-thick) were collected and contrasted with 1% aqueous uranyl acetate and 0.3% Reynold’s lead citrate to assess the ultrastructural preservation of the sample. Protein A (ProtA, Cell Microscopy Center, Utrecht, The Netherlands) coupled to 10 nm gold particles were applied to semithin sections to serve as fiducial markers for tomographic reconstructions.

##### Electron Tomography and Data Analysis

MF synapses were identified by their distinct morphology (large boutons, multiple AZs) and target specificity (primary dendrites of CA3 PCs) (Chicurel and Harris, 1992). Synapses were selected for tomography when the PSD was juxtaposed to a cluster of SVs in the presynaptic compartment and the synaptic cleft was parallel to the tilt axis and clearly visible at 0° stage tilt. Only MF-CA3 PC spine synapses were included in the analysis. Glutamatergic SC spine synapses were identified based on their location within the CA1 region of the hippocampal slice and according to well-established ultrastructural features such as the presence of a small postsynaptic compartment lacking mitochondria or microtubules (Imig et al., 2014).

Single-axis tilt series were acquired on a JEOL JEM-2100 200kV transmission electron microscope from −60° to + 60° in 1° increments and binned by a factor of two at 30,000 times magnifications with an Orius SC1000 camera (Gatan) using SerialEM for acquiring automated tilt series (Mastronarde, 2005). Tomograms were reconstructed and binned by a factor of three (1.554 nm isotropic voxel size of the final tomogram) and segmented for analysis using the IMOD package (Kremer et al., 1996). All vesicles were segmented manually as perfect spheres with the center being placed into the tomographic slice in which the vesicle diameter appeared to be the largest (i.e. when the vesicle is cut at its midline). The sphere outline was adjusted to lie on the center of the outer leaflet of the vesicle lipid bilayer. Non-spherical organelles (e.g. endoplamisc reticulum, tubular endosomal intermediates) were occasionally observed in tomographic reconstructions and excluded from the analysis. The active zone (AZ) was defined as the region of presynaptic membrane that was apposed to the postsynaptic density (PSD). In cryopreserved tissue, PSDs often appeared less electron-dense than in chemically fixed material (see for example Figure S1). Therefore we used the widening of the synaptic cleft as a second morphological feature on which to base the AZ area for segmentation. The AZ was segmented at the center of the inner leaflet of the presynaptic lipid bilayer.

In segmented models, the shortest distance between vesicle membranes and the AZ were calculated using the mtk program of the IMOD package (Kremer et al., 1996). Vesicle diameters and AZ surface areas were extracted from segmented models using the imodinfo program. A vesicle was defined as docked when there was no measurable distance between the outer leaflet of the vesicle and the inner leaflet of the lipid bilayer (i.e. when the dark pixels corresponding to the vesicular outer leaflet were contiguous with those of the inner plasma membrane leaflet). Based on the voxel size of 1.554 nm, these ‘docked’ vesicles fall into the 0-2 nm bin. Number of vesicles in discrete bins (i.e. 0-2 nm, 0-40 nm, and 0-100 nm from the AZ membrane) were normalized to the measured AZ area and reported as number of vesicles per 0.01 µm^2^ AZ. To allow a more direct comparison of our results to data obtained from 2D-analyses of SV docking (Chang et al., 2018), we also report the number of SVs within 5 nm of the AZ membrane for each condition analyzed. Vesicles with a diameter less than 60 nm were classified as SVs in the analysis, whereas vesicles with a diameter exceeding 60 nm and having no prominent dense core were classified as giant vesicles (GVs). The 60 nm cutoff was chosen, because it exceeded three standard deviations from the mean SV diameter measured in SC synapses (mean: 43.44 nm, standard deviation: 3.92 nm). Vesicles that contained a prominent electron-dense core were considered dense-cored vesicles (DCVs). For illustrative purposes, figures depicting tomographic sub-volumes represent an overlay of seven consecutive tomographic slices (10.88 nm-thick sub-volume) unless otherwise specified and were generated using the IMOD package (Kremer et al., 1996).

##### RRP Calculations

In MF-CA3 synapses, calculations of mean docked vesicle numbers per AZ and per MFB were based on our estimates of the mean number of docked vesicles per unit AZ area (0.9 SVs, 0.1 GVs and 0.04 DCVs per 0.01 µm^2^), and previously published estimates of the mean MF AZ surface area (0.12 µm^2^) and mean AZ number (29.75 AZs) per MFB in P28 rat MFBs (Rollenhagen et al., 2007). Our calculations of total docked vesicle numbers per MFB neglect filopodial extensions. We calculated the mean docked vesicle numbers per AZ (10.7 SVs, 1.2 GVs, 0.5 DCV) and per MFB (∼320 SVs, 37 GVs, 15 DCVs) in mature (DIV28) MF synapses as well as the mean number of total membrane-proximal (within 0-40 nm of the AZ; 24.6 SVs, 4 GVs, 2.6 DCVs) vesicles per AZ. ET further enabled precise volume and membrane surface area measurements for all docked vesicles of a given morphological type (mean Ø; SV, 45.17 nm; GV, 85.77 nm; DCV 74.41 nm). We further assumed a specific membrane capacitance of 1 μF/cm^2^ (Hallermann et al., 2003) to determine that the fusion of all docked vesicles, irrespective of their type, would change the membrane capacitance (ΔC_m_) by ∼32 fF per MFB and that the fusion of all docked and tethered vesicles together would correspond to a change in ΔC_m_ by ∼88 fF.

##### Electrophysiology

All recordings from CA3 PCs in organotypic slice cultures were performed at DIV14. Prior to recordings, slices were incubated for 30 min in an interface chamber with carbogen-saturated ACSF (120 mM NaCl, 26 mM NaHCO_3_, 10 mM D-glucose, 2 mM KCl, 2 mM MgCl_2_, and 2 mM CaCl_2_, and 1 mM KH_2_PO_4_ - 304 mOsm). One or two CA3 PCs were then whole-cell voltage clamped using a glass pipette (2.5-3.0 MΩ) filled with internal solution (100 mM KCl, 50 mM K-gluconate, 10 mM HEPES, 4 mM ATP-Mg, 0.3 mM GTP-Na, 0.1 mM EGTA, and 0.4% biocytin, pH 7.4, 300 mOsm) and the holding potential was set at −70 mV using an EPC-10 amplifier [Patchmaster 2 software (HEKA/Harvard Bioscience)]. For measurements of mEPSC amplitudes and frequencies, slices were initially perfused with 1 µM TTX and 10 µM bicuculline and mEPSCs were then recorded for 10 min, after which the slices were perfused with 1 µM TTX, 10 µM bicuculline, and 2 µM DCG-IV for 15 min to record DCG-IV insensitive mEPSCs. Measurements of all mEPSCs (TTX/bicuculline) were recorded in two-5 min epochs, while measurements of non-MF mEPSCs (TTX/bicuculline/DCG-IV) were recorded in three-5 min epochs. The last epochs of each recording were used for mEPSC analysis. All electrophysiological traces were analyzed using Axograph X software (AxoGraph Scientific) using a template fit algorithm for automatic event detection (Jonas et al., 1993; Pernía-Andrade et al., 2012). After recordings, some slices were fixed and biocytin-filled CA3 PC were stained with Alexa Fluoro-555-labeled streptavidin (see Light Microscopic Analysis section for detailed procedure).

Only cells that had a reduction in the mEPSC frequency upon application of DCG-IV were included in the analysis and the threshold for mEPSC detection was set to 8 pA. The number of events with a given mEPSC amplitude after DCG-IV application (1 pA bins) was subtracted from the number of events prior to DCG-IV application assuming the events lost upon application of DCG-IV were all of MF origin. The vesicle diameter was measured from the edge of the outer leaflet of the vesicle lipid bilayer. To account for the volume of the vesicle lumen, the lipid bilayer (approx. 4 nm-thick as measured from center-to-center of the vesicle bilayer) subtracted from the diameter size (8 nm total) and the volume of the vesicle lumen was calculated. Based on these assumptions, we calculated that an mEPSC amplitude of approximately 30 pA would arise from a quanta released from a vesicle with a diameter of approximately 60 nm.

##### Light Microscopic Analysis

To demonstrate the correct anatomical organization of the MF pathway in organotypic slices, slices were removed from culture inserts and fixed by overnight immersion in 4% PFA in 0.1 M PB (pH 7.4). Slices were washed in 0.1 M PB (pH 7.4) and then incubated overnight at 4°C in 10% normal goat serum (NGS), 0.3% Triton X-100, and 0.1% cold water fish skin gelatin (FSG) in 0.1 M PB (pH 7.4) to permeabilize membranes and block non-specific binding sites. Slices were incubated overnight at 4°C in 5% NGS, 0.3% Triton X-100 and 0.1% FSG in 0.1 M PB (pH 7.4) containing primary antibodies against SV clusters within the synaptic terminals of MF projections [polyclonal rabbit anti-Synaptoporin, SYnaptic SYstems (Cat# 102 003), 1:1000 dilution] and cell bodies and dendritic arborizations [polyclonal chicken anti-MAP2, Novus Biologicals (Cat# NB300-213), 1:600 dilution]. Slices were washed in 0.1 M PB (pH 7.4) and primary antibodies were visualized by a 2 hr incubation at RT in 5% NGS, 0.1% Triton X-100 and 0.1% FSG in 0.1 M PB (pH 7.4) containing goat anti–rabbit Alexa 555 [Thermo Fisher (Cat# A21429), dilution 1:1000] and goat anti-chicken Alexa 488 [Thermo Fisher(Cat# A-11039), dilution 1:1000]. Following final washing steps in 0.1 M PB (pH 7.4), slices were floated onto Superfrost™ glass slides with the membrane confetti in contact with the slide and Menzel-Gläser #1.5 glass coverslips were mounted using Aqua-Poly/Mount mounting medium (Polysciences, Inc., Cat# 18606-20).

To localize active zone release sites within mossy fiber boutons, slices were removed from culture inserts and fixed by overnight immersion in 4% PFA in 0.1 M PB (pH 7.4). Slices were washed in 0.1 M PB (pH 7.4) and then cryoprotected by immersion in an increasing gradient of 10%, 20%, and 30% sucrose in 0.1 M PB (pH 7.4) until saturation. Slices were placed flat, slice-side down (confetti-side up), on the inner base of a quadratic 10 × 10 × 10 mm form made out of aluminium foil and carefully covered with liquid Tissue-Tek^®^ OCT compound (Sakura, Cat# 4583) without introducing air bubbles. The OCT-filled form was then rapidly frozen on a liquid nitrogen-cooled aluminium block, the foil was removed and the frozen OCT block was mounted slice-side up on a specimen stub with OCT in a precooled (specimen holder, −18°C; chamber, −18°C) cryostat (Leica CM3050 S). Once the temperature of the embedded slice had equilibrated, unnecessary OCT compound was trimmed away with a razor blade and 10 µm-thick cryosections were made through the organotypic slice with the aid of a glass anti-roll plate and thaw-mounted on Superfrost™ slides. Slides were air-dried at RT for 30 min and a hydrophobic pen (DAKO, Cat# S2002) was used to delineate the border of the slide surface. Slides were washed briefly in 0.1 M PB (pH 7.4) and incubated 90 min at RT in 10% NGS, 0.3% Triton X-100, and 0.1% FSG in 0.1 M PB (pH 7.4). Slices were then incubated overnight at 4°C in 3% NGS, 0.3% Triton X-100 and 0.1% FSG in 0.1 M PB (pH 7.4) containing primary antibodies for the detection of SV clusters within the synaptic terminals of MF projections [polyclonal rabbit anti-Synaptoporin, SYnaptic SYstems (Cat# 102 003), 1:1000 dilution] and presynaptic active zones [monoclonal mouse anti-Bassoon, Enzo Life Sciences (Cat# SAP7F407), 1:400 dilution]. Slices were washed in 0.1 M PB (pH 7.4) and primary antibodies were visualized by 2 hr incubation at RT in 5% NGS, 0.1% Triton X-100 and 0.1% FSG in 0.1 M PB (pH 7.4) containing goat anti-rabbit Alexa 488 [Thermo Fisher (Cat# A11008), dilution 1:1000] and goat anti-mouse Alexa 555 [Thermo Fisher (Cat# A21424), dilution 1:1000]. Following a brief wash in 0.1 M PB, slides were dipped in distilled water and Menzel-Gläser #1,5 coverslips were mounted using Aqua-Poly/Mount mounting medium (Polysciences, Inc., Cat# 18606-20).

Confocal light microscopic analysis of biocytin-filled CA3 pyramidal cells was performed to validate that electrophysiological recordings were of the correct, as assessed by the anatomical location within the hippocampal subfields and morphological features (i.e, pyramidal soma, presence of large, complex spines in the proximal regions of apical dendritic arborizations) of the filled cell (see Figure S1D). Immediately following mEPSC recordings and removal of the patch pipette, biocytin-filled CA3 pyramidal cells (see Electrophysiology section above for detailed procedure) were fixed for light microscopic analysis by overnight immersion of the slice in 4% PFA in 0.1 M PB (pH 7.4). Slices were washed in 0.1 M PB (pH 7.4) and then incubated overnight at 4°C in 10% NGS, 0.3% Triton X-100, and 0.1% FSG in 0.1 M PB (pH 7.4). Biocytin-filled cells were visualized by incubation of slices for 3 hrs at RT in Steptavidin-Alexa 555 [1:500 dilution] in 5% NGS, 0.1% Triton X-100 and 0.1% FSG in 0.1 M PB (pH 7.4). Slices were washed in 0.1 M PB (pH 7.4) and cell nuclei were labeled by a 30 min incubation in DAPI [300 nM in 0.1 M PB]. Following final washing steps in 0.1 M PB (pH 7.4), slices were floated onto Superfrost glass slides with the membrane confetti in contact with the slide and Menzel-Gläser #1,5 glass coverslips were mounted using Aqua-Poly/Mount mounting medium (Polysciences, Inc., Cat# 18606-20).

Confocal laser scanning micrographs of were acquired with a Leica TCS-SP5 confocal microscope equipped with a tunable white light laser, a resonant scanner, hybrid GaAsP detectors, and a motorized stage. Tiled z-series were acquired with (i) a HCX PL APO 40.0x (NA=1.25) oil immersion objective to generate low magnification overviews of entire organotypic slices (pinhole = 3.0 AU, voxel size x, y, z = 0.3, 0.3, 2 µm)(Figure S1A) and reconstructions of biocytin-filled pyramidal neurons within CA3 *stratum pyramidale* (pinhole = 1.0 AU, voxel size x, y, z = 95, 95, 335 nm)(Figure S1D), or with (ii) a HCX PL APO CS 100x (NA=1.4) oil immersion objective to visualize mossy fiber terminals within CA3 *stratum lucidum* (pinhole = 1.0 AU, voxel size x, y, z = 89, 89, 130 nm) and high magnification reconstructions of complex postsynaptic spines (thorny excrescences) emerging from the proximal dendrites of biocytin-filled CA3 pyramidal neurons (pinhole = 0.5 AU, voxel size x, y, z = 47, 47, 130 nm). For illustration purposes thorny excrescences were subjected to spatial deconvolution by use of two ImageJ (National Institutes of Health; Bethesda, MD) plugins: point spread functions (PSF) were generated using Diffraction PSF 3D plugin and iterative deconvolution was performed with the Richardson-Lucy algorithm (DeconvolutionLab plugin; Biomedical Imaging Group, EPFL; Lausanne, Switzerland).

#### Quantification and Statistical Analysis

Data are represented as mean ± SEM unless indicated otherwise. Statistical analyses were carried out using GraphPad Prism software 7 (* when P<0.05; ** when P<0.01, and *** when P<0.001). For comparisons of two conditions (i.e. SC and MF synaptic profiles from DIV14 WT slice cultures; Fig 2A-G) statistical difference were determined by an unpaired t-test when the data set was normally distributed as determined by a KS normality test, and by a Mann-Whitney unpaired t-test if it was not normally distributed. For pharmacological manipulation experiments, statistical significance was tested by one-way ANOVA with Bonferroni correction as a post-test if the data set was normally distributed. If the data set was not normally distributed, Kruskal-Wallis ANOVA test with a Dunn’s comparison of all columns was performed as a post-test to probe for statistical significance. The number of tomograms analyzed for each experiment (n), the number of slice cultures or animals used (N), and all EM data is summarized in Table S1. Statistics were performed based on the number of tomograms for each sample with the exception of docked vesicle diameters and unattached GVs and DCVs. In the latter scenarios, the number of vesicles was used for statistical analysis and is noted in parentheses. Electrophysiological recordings were performed on 28 cells from two independent WT slice cultures at DIV14. Statistical difference for electrophysiological experiments was determined by Wilcoxon matched pairs signed rank tests.

---

**Figure S1.**
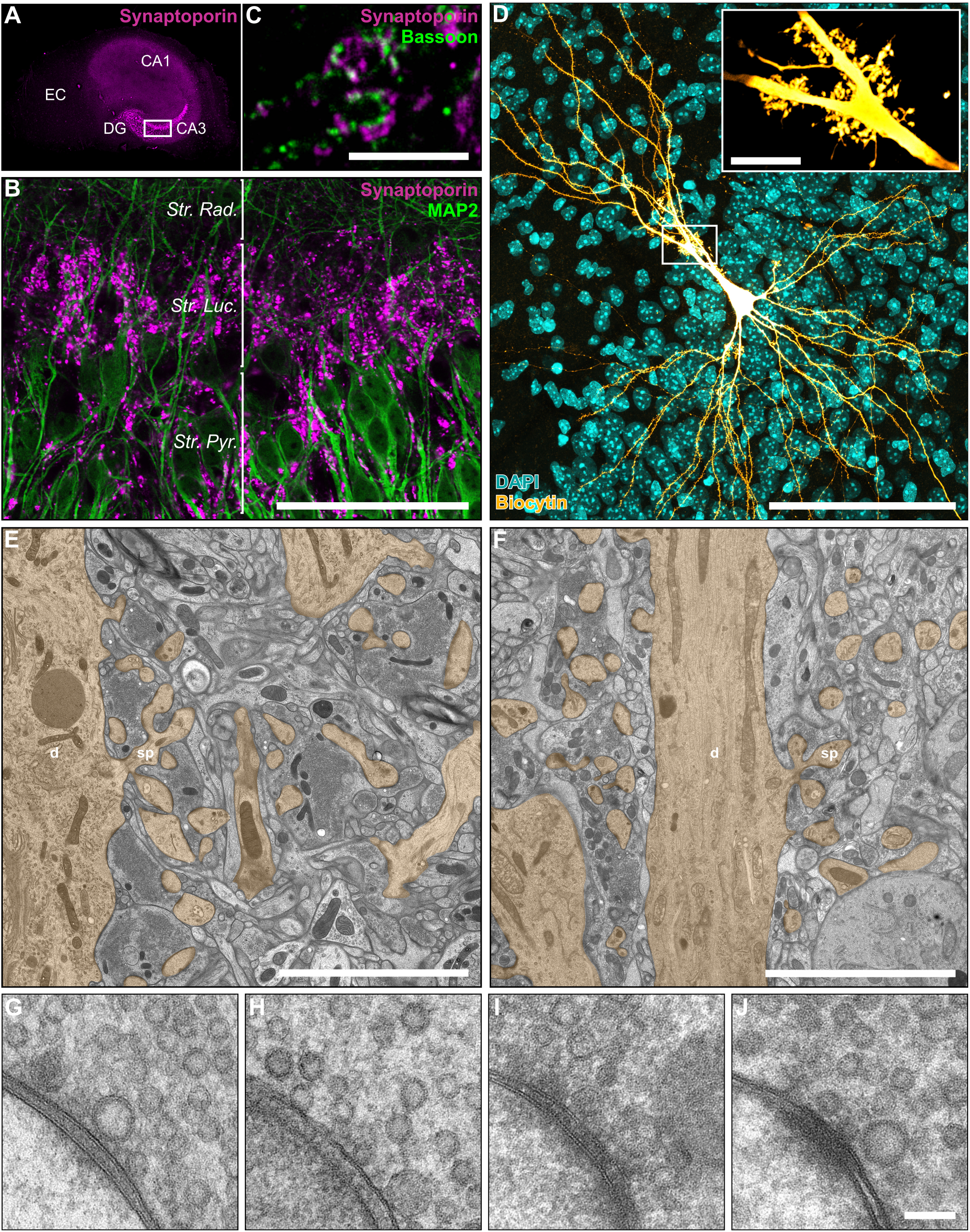
Morphological Characterization of the MF-CA3 PC Synapse, Related to Figure 1. (**A**-**F**) Organotypic organization of the MF-CA3 connection is preserved in cultured hippocampal slices. (**A**) Immunoreactivity for the MFB SV protein Synaptoporin (magenta) is restricted to the CA3 *stratum lucidum*. (**B**) Synaptoporin-positive MFBs (magenta) contact MAP2-immunoreactive primary dendrites of CA3 PCs (green) within the *stratum lucidum*. (**C**) Synaptoporin-positive MFBs (magenta) exhibit multiple AZs as indicated by the presence of Bassoon-positive cluster (green). (**D**) A biocytin-filled CA3 PC (’fire’ lookup table) in a cultured hippocampal slice is embedded in the CA3 PC layer (DAPI, cyan) and exhibits several complex, multi-headed spines (thorny excrescences; insert). (**E**, **F**) Electron micrograph of the *stratum lucidum* from a HPF cultured slice (**E**) and a perfusion-fixed hippocampus (**F**) [postsynaptic elements including dendrites (d) and spines (sp) in orange]. (**G**-**J**) Electron micrographs of MF-CA3 PC spine synapses from a cultured hippocampal slice prepared by HPF (DIV28; **G**), from an acute brain slice prepared at postnatal day (P)18 by HPF (**H**), and from perfusion fixed tissue (P28) with either 4% PFA, 2.5% GA in 0.1M PB (Fixative 1; **I**) or 2% PFA, 2.5% GA in 0.1 cacodylate buffer (Fixative 2; **J**). Abbreviations: EC, entorhinal cortex; DG, dentate gyrus; CA1/3, *Cornu Ammonis* 1 and 3; Str. Rad., *Stratum Radiatum*; Luc., *Lucidum*; Pyr., *Pyramidale*; d, dendrite; sp, spine. Scale bars: C, E, F, 5 µm; B and D, 100 µm; insert in D, 10 µm; G-J, 100 nm.

**Figure S2.**
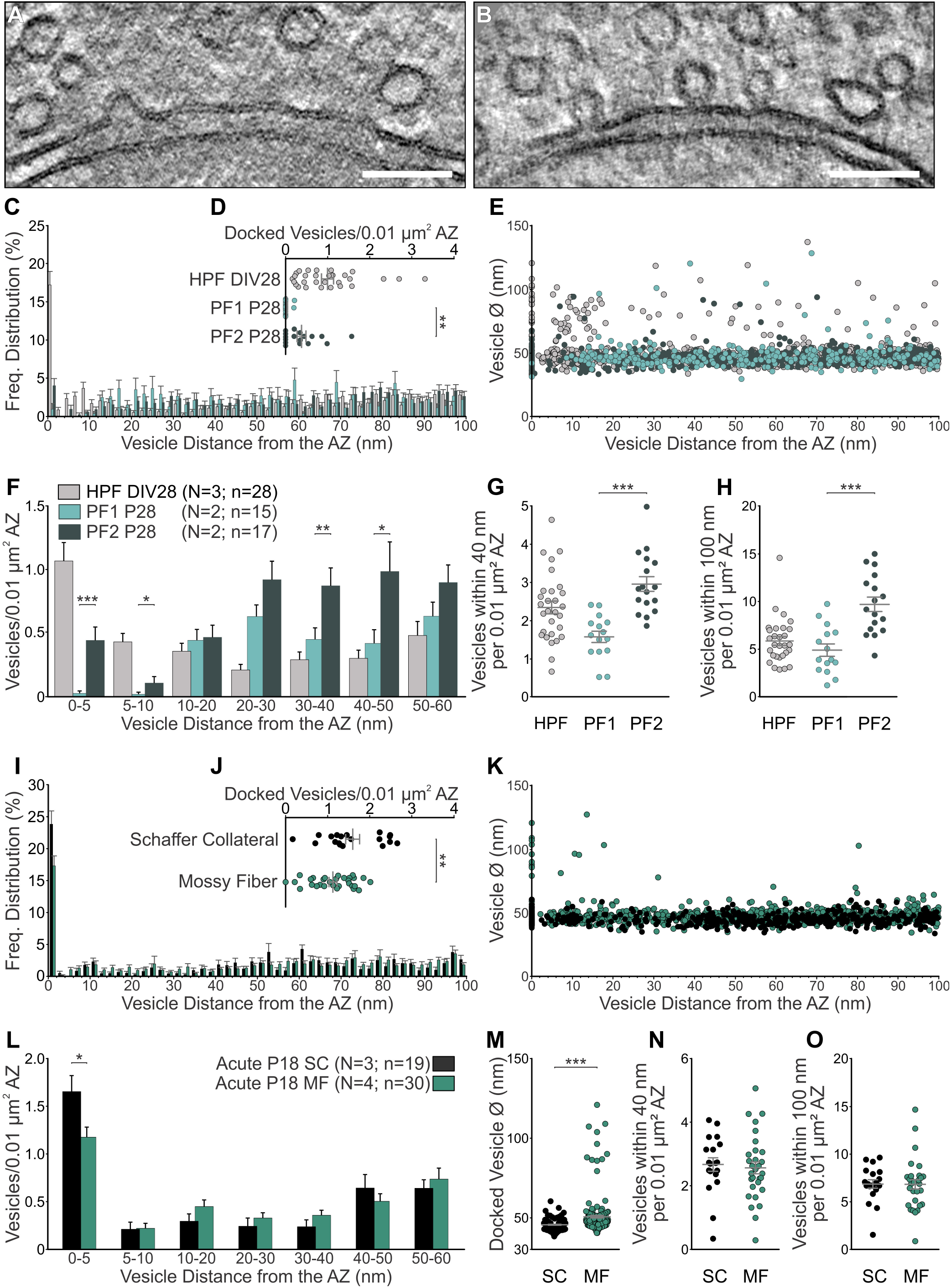
Ultrastructural Effects of Different Sample Preparation and Fixation Methods on Presynaptic SV Pools, Related to Figure 2. (**A**-**H**) Comparison of MF synaptic ultrastructure in postnatal day (P) 28 WT mice perfused with either 4% paraformaldehyde (PFA), 2.5% glutaraldehyde (GA) in 0.1 M phosphate buffer (Fixative 1; ET subvolume shown in **A**) or with 2% PFA, 2.5% GA, 2 mM CaCl_2_ in 0.1 M cacodylate buffer (Fixative 2; ET subvolume shown in **B**) with synaptic morphology of WT slice cultures cryo-fixed at DIV28. See Table S1D (**C**) Spatial distribution of vesicles within 100 nm of the AZ membrane in perfusion-fixed material from P28 WT mice (N = 2 mice; Fixative 1 n = 15 tomograms; Fixative 2 n = 17) and WT slice cultures cryo-fixed at DIV28 (N = 3 cultures; n = 28). (**D**) Scatterplot of docked vesicles normalized to AZ area from chemically fixed and age-matched slice cultures. (**E**) Plot of vesicle diameters for all vesicles analyzed and their respective distance to the AZ membrane. (**F**) Mean number of vesicles within bins of 5 nm and 10 nm from the AZ normalized to AZ area. (**G**, **H**) Scatterplots of vesicles within 40 nm (**G**) and 100 nm (**H**) of the AZ membrane normalized to AZ area. (**I**-**O**) Analysis of vesicle pools in SC (N = 3 mice; n =19 tomograms) and MF synapses (N = 4 mice; n = 30 tomograms) in acute brain slices prepared at P18 by HPF (frozen in 20% BSA in ACSF) immediately after dissection. See Table S1E (**I**) Spatial distribution of vesicles within 100 nm of the active zone (AZ) membrane in SC and MF synapses. (**J**) Scatterplots of the mean number of docked vesicles normalized to AZ area. (**K**) Plot of vesicle diameters for all vesicles analyzed and their respective distance to the AZ membrane. (**L**) Mean number of vesicles within bins of 5 nm and 10 nm from the AZ normalized to AZ area. (**M**) Scatterplot of vesicle diameters for all docked vesicles analyzed in SC (n = 113) and MF (n = 197) synapses. (**N**, **O**) Scatterplots of vesicles within 40 nm (**N**) and 100 nm (**O**) of the AZ membrane normalized to AZ area. Scale bars: A and B, 100 nm. Values indicate mean ± SEM; *p<0.05; **p<0.01; ***p<0.001.

**Figure S3.**
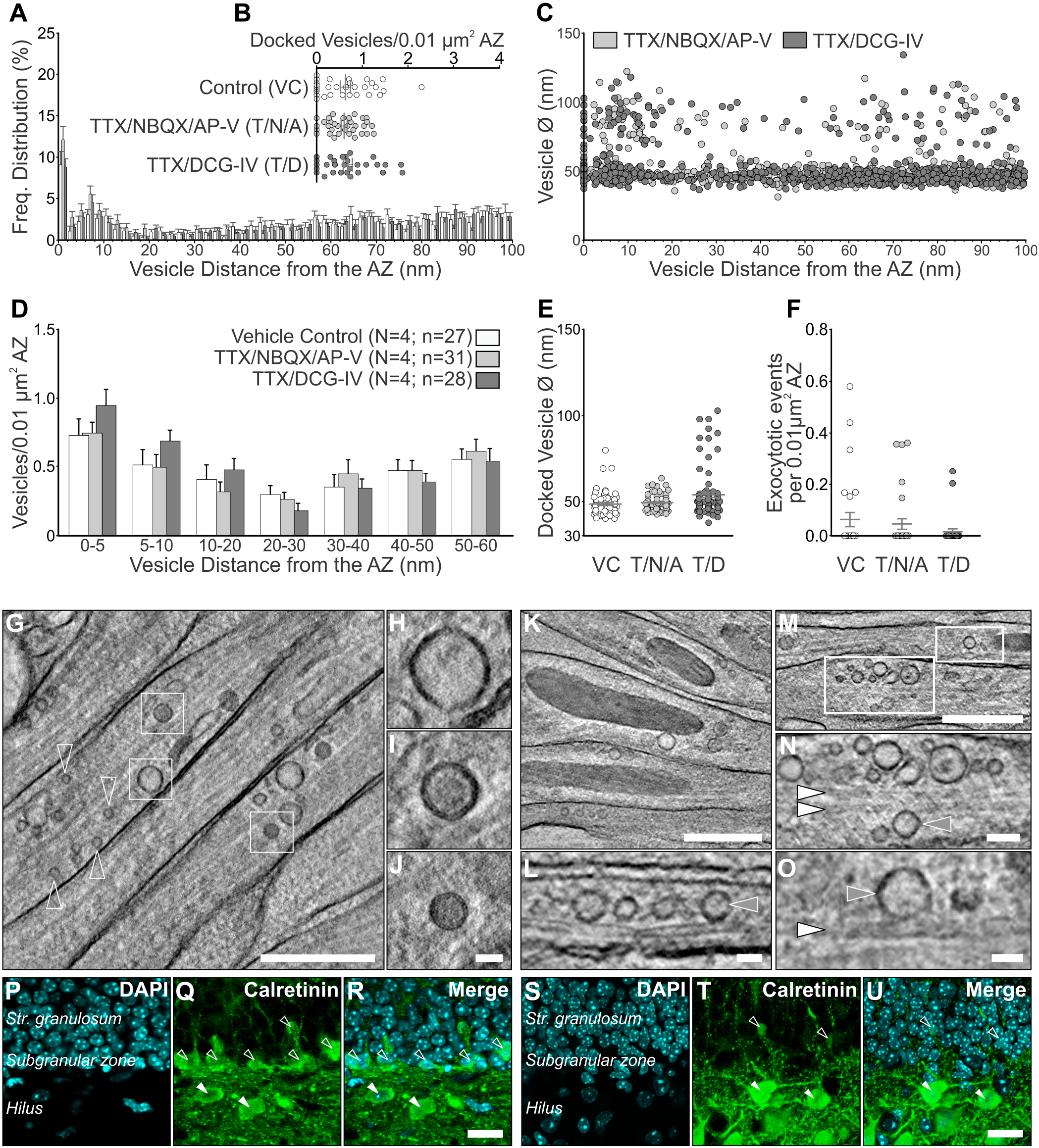
Investigating the Origin of GVs in MFBs, Related to Figure 3. (**A**-**F**) Characterization of vesicle pools in MF synapses (N = 4 cultures) treated for 10 min with either vehicle control (VC; slice culture medium; n = 27 tomograms), 1 µM TTX, 2 µM NBQX, and 50 µm AP-V (T/N/A; n = 31), or 1 µM TTX and 2 µM DCG-IV (T/D; n = 28). See Table S1F (**A**) Spatial distribution of vesicles within 100 nm of the AZ membrane in MF synapses. (**B**) Scatterplot of docked vesicles normalized to AZ area. (**C**) Plot of vesicle diameters for all vesicles analyzed and their respective distance to the AZ membrane. (**D**) Mean number of vesicles within bins of 5 and 10 nm from the AZ normalized to AZ area. (**E**) Scatterplot of vesicle diameters for all docked vesicles analyzed (VC = 62; T/N/A = 72; T/D = 83 vesicles) (**F**) Scatterplot of the number of morphological exocytotic events per tomogram normalized to AZ area. (**G**-**O**) Morphological characterization of vesicle pools in MF axon bundles in the *stratum lucidum* of P18 acute hippocampal slices. (**G**, **K**, and **M**) Tomographic subvolume (42 nm-thick) through MF axon bundles. White boxes indicate regions enlarged in H-J, L, N, and O. (**H**-**J**) Mossy fiber axons contain multiple vesicle classes, including small clear-cored vesicles (open arrowheads), large clear-cored vesicles (**H**), and DCVs (**I** and **J**) (**L**, **N**, and **O**) Single tomographic slices (2.8 nm-thick) reveal close contact between large clear-cored vesicles (grey arrowheads; **L**, Ø=60 and 64 nm; **N**, Ø=87 nm; **O**, Ø=81 nm) and microtubules (white arrowheads), indicative of active axonal transport between granule cell bodies and MFBs. (**P**-**U**) Characterization of neurogenesis in the hippocampus of P28 mice (**P**-**R**) and cultured hippocampal slices at DIV28 (**S**-**U**). Adult mice exhibit calretinin-positive (green) immature granule cells (open arrow head) in the subgranular zone of the dentate gyrus (cell bodies in cyan labeled by DAPI) and hilar Mossy cells (white arrowhead). In cultured hippocampal slices, calretinin-immunoreactivity is almost exclusively restricted to hilar Mossy cells, indicating an almost complete loss of immature granule cells in the DG. Scale bars: G, K, and M, 500 nm; N, 100 nm; H-J, L, O, 50 nm, and R, U, 20 µm Values indicate mean ± SEM; *p<0.05; **p<0.01; ***p<0.001.

## REFERENCES

Amaral, D.G., and Dent, J.A. (1981). Development of the mossy fibers of the dentate gyrus: I. A light and electron microscopic study of the mossy fibers and their expansions. J. Comp. Neurol. 195, 51–86.

Augustin, I., Rosenmund, C., Südhof, T.C., and Brose, N. (1999). Munc13-1 is essential for fusion competence of glutamatergic synaptic vesicles. Nature 400, 457–461.

Battistin, T., and Cherubini, E. (1994). Developmental Shift From Long□term Depression to Long-term Potentiation at the Mossy Fibre Synapses in the Rat Hippocampus. Eur. J. Neurosci. 6, 1750–1755.

Blatow, M., Caputi, A., Burnashev, N., Monyer, H., and Rozov, A. (2003). Ca2+ Buffer Saturation Underlies Paired Pulse Facilitation in Calbindin-D28k-Containing Terminals. Neuron 38, 79–88.

Brandt, M.D., Jessberger, S., Steiner, B., Kronenberg, G., Reuter, K., Bick-Sander, A., Behrens, W. von der, and Kempermann, G. (2003). Transient calretinin expression defines early postmitotic step of neuronal differentiation in adult hippocampal neurogenesis of mice. Mol. Cell. Neurosci. 24, 603–613.

Chang, S., Trimbuch, T., and Rosenmund, C. (2018). Synaptotagmin-1 drives synchronous Ca2+-triggered fusion by C2B-domain-mediated synaptic-vesicle-membrane attachment. Nat. Neurosci. 21, 33–42.

Chicurel, M.E., and Harris, K.M. (1992). Three-dimensional analysis of the structure and composition of CA3 branched dendritic spines and their synaptic relationships with mossy fiber boutons in the rat hippocampus. J. Comp. Neurol. 325, 169–182.

Cho, R.W., Buhl, L.K., Volfson, D., Tran, A., Li, F., Akbergenova, Y., and Littleton, J.T. (2015). Phosphorylation of Complexin by PKA Regulates Activity-Dependent Spontaneous Neurotransmitter Release and Structural Synaptic Plasticity. Neuron 88, 749–761.

Dittman, J., and Ryan, T.A. (2009). Molecular circuitry of endocytosis at nerve terminals. Annu. Rev. Cell Dev. Biol. 25, 133–160.

Éltes, T., Kirizs, T., Nusser, Z., and Holderith, N. (2017). Target Cell Type-Dependent Differences in Ca2+ Channel Function Underlie Distinct Release Probabilities at Hippocampal Glutamatergic Terminals. J. Neurosci. 37, 1910–1924.

Galimberti, I., Gogolla, N., Alberi, S., Santos, A.F., Muller, D., and Caroni, P. (2006). Long-Term Rearrangements of Hippocampal Mossy Fiber Terminal Connectivity in the Adult Regulated by Experience. Neuron 50, 749–763.

Geiger, J.R.P., and Jonas, P. (2000). Dynamic Control of Presynaptic Ca2+ Inflow by Fast-Inactivating K+ Channels in Hippocampal Mossy Fiber Boutons. Neuron 28, 927–939.

Hallermann, S., Pawlu, C., Jonas, P., and Heckmann, M. (2003). A large pool of releasable vesicles in a cortical glutamatergic synapse. Proc. Natl. Acad. Sci. 100, 8975–8980.

Hayat, M.A. (1981). Principles and techniques of electron microscopy. Biological applications. (Cambridge University Press).

He, E., Wierda, K., van Westen, R., Broeke, J.H., Toonen, R.F., Cornelisse, L.N., and Verhage, M. (2017). Munc13-1 and Munc18-1 together prevent NSF-dependent de-priming of synaptic vesicles. Nat. Commun. 8, 15915.

He, L., Xue, L., Xu, J., McNeil, B.D., Bai, L., Melicoff, E., Adachi, R., and Wu, L.-G. (2009). Compound vesicle fusion increases quantal size and potentiates synaptic transmission. Nature 459, 93–97.

Helassa, N., Dürst, C.D., Coates, C., Kerruth, S., Arif, U., Schulze, C., Wiegert, J.S., Geeves, M., Oertner, T.G., and Török, K. (2018). Ultrafast glutamate sensors resolve high-frequency release at Schaffer collateral synapses. Proc. Natl. Acad. Sci. U. S. A. 115, 5594–5599.

Henze, D.A., Card, J.P., Barrionuevo, G., and Ben-Ari, Y. (1997). Large Amplitude Miniature Excitatory Postsynaptic Currents in Hippocampal CA3 Pyramidal Neurons Are of Mossy Fiber Origin. J. Neurophysiol. 77, 1075–1086.

Henze, D.A., McMahon, D.B.T., Harris, K.M., and Barrionuevo, G. (2002). Giant Miniature EPSCs at the Hippocampal Mossy Fiber to CA3 Pyramidal Cell Synapse Are Monoquantal. J. Neurophysiol. 87, 15–29.

Imig, C., and Cooper, B.H. (2017). 3D analysis of synaptic ultrastructure in organotypic hippocampal slice culture by high-pressure freezing and electron tomography. In Methods in Molecular Biology, pp. 215–231.

Imig, C., Min, S.-W., Krinner, S., Arancillo, M., Rosenmund, C., Südhof, T.C.C., Rhee, J., Brose, N., and Cooper, B.H.H. (2014). The Morphological and Molecular Nature of Synaptic Vesicle Priming at Presynaptic Active Zones. Neuron 84, 416–431.

Jackman, S.L., Turecek, J., Belinsky, J.E., and Regehr, W.G. (2016). The calcium sensor synaptotagmin 7 is required for synaptic facilitation. Nature 529, 88–91.

Jonas, P., Major, G., and Sakmann, B. (1993). Quantal components of unitary EPSCs at the mossy fibre synapse on CA3 pyramidal cells of rat hippocampus. J. Physiol. 472, 615–663.

Kaeser, P.S., and Regehr, W.G. (2017). The readily releasable pool of synaptic vesicles. Curr. Opin. Neurobiol. 43, 63–70.

Kamiya, H., and Yamamoto, C. (1997). Phorbol ester and forskolin suppress the presynaptic inhibitory action of group-II metabotropic glutamate receptor at rat hippocampal mossy fibre synapse. Neuroscience 80, 89–94.

Kamiya, H., Shinozaki, H., and Yamamoto, C. (1996). Activation of metabotropic glutamate receptor type 2/3 suppresses transmission at rat hippocampal mossy fibre synapses. J. Physiol. 493, 447–455.

Korogod, N., Petersen, C.C., and Knott, G.W. (2015). Ultrastructural analysis of adult mouse neocortex comparing aldehyde perfusion with cryo fixation. Elife 4, e05793.

Kremer, J.R., Mastronarde, D.N., and McIntosh, J.R.R. (1996). Computer visualization of three-dimensional image data using IMOD. J. Struct. Biol. 116, 71–76.

Laatsch, R.H., and Cowan, W.M. (1966). Electron microscopic studies of the dentate gyrus of the rat. I. Normal structure with special reference to synaptic organization. J. Comp. Neurol. 128, 359–395.

Lawrence, J.J., Grinspan, Z.M., and McBain, C.J. (2004). Quantal transmission at mossy fibre targets in the CA3 region of the rat hippocampus. J. Physiol. 554, 175–193.

Lee, J.S., Ho, W.-K., and Lee, S.-H. (2012). Actin-dependent rapid recruitment of reluctant synaptic vesicles into a fast-releasing vesicle pool. Proc. Natl. Acad. Sci. 109, E765–E774.

Marchal, C., and Mulle, C. (2004). Postnatal maturation of mossy fibre excitatory transmission in mouse CA3 pyramidal cells: A potential role for kainate receptors. J. Physiol. 561, 27–37.

Mastronarde, D.N. (2005). Automated electron microscope tomography using robust prediction of specimen movements. J. Struct. Biol. 152, 36–51.

Midorikawa, M., and Sakaba, T. (2017). Kinetics of Releasable Synaptic Vesicles and Their Plastic Changes at Hippocampal Mossy Fiber Report Kinetics of Releasable Synaptic Vesicles and Their Plastic Changes at Hippocampal Mossy Fiber Synapses. Neuron 96, 1033–1040.e3.

Miki, T., Nakamura, Y., Malagon, G., Neher, E., and Marty, A. (2018). Two-component latency distributions indicate two-step vesicular release at simple glutamatergic synapses. Nat. Commun. 9, 3943.

Mori, M., Abegg, M.H., Gähwiler, B.H., and Gerber, U. (2004). A frequency-dependent switch from inhibition to excitation in a hippocampal unitary circuit. Nature 431, 453–456.

Neher, E. (2015). Merits and Limitations of Vesicle Pool Models in View of Heterogeneous Populations of Synaptic Vesicles. Neuron 87, 1131–1142.

Neher, E., and Brose, N. (2018). Dynamically Primed Synaptic Vesicle States: Key to Understand Synaptic Short-Term Plasticity. Neuron 100, 1283–1291.

Nicoll, R.A., and Schmitz, D. (2005). Synaptic plasticity at hippocampal mossy fibre synapses. Nat. Rev. Neurosci. 6, 863–876.

Oertner, T.G., Sabatini, B.L., Nimchinsky, E.A., and Svoboda, K. (2002). Facilitation at single synapses probed with optical quantal analysis. Nat. Neurosci. 5, 657–664.

Pernía-Andrade, A.J., Goswami, S.P., Stickler, Y., Fröbe, U., Schlögl, A., and Jonas, P. (2012). A Deconvolution-Based Method with High Sensitivity and Temporal Resolution for Detection of Spontaneous Synaptic Currents In Vitro and In Vivo. Biophys. J. 103, 1429–1439.

Raineteau, O., Rietschin, L., Gradwohl, G., Guillemot, F., and Gähwiler, B.H. (2004). Neurogenesis in hippocampal slice cultures. Mol. Cell. Neurosci. 26, 241–250.

Regehr, W.G. (2012). Short-term presynaptic plasticity. Cold Spring Harb. Perspect. Biol. 4, a005702.

Rollenhagen, A., Sätzler, K., Patricia Rodríguez, E., Jonas, P., Frotscher, M., Lübke, J.H.R., Satzler, K., Rodriguez, E.P., Jonas, P., Frotscher, M., et al. (2007). Structural Determinants of Transmission at Large Hippocampal Mossy Fiber Synapses. J. Neurosci. 27, 10434–10444.

Rosenmund, C., and Stevens, C.F. (1997). The rate of aldehyde fixation of the exocytotic machinery in ultured hippocampal synapses. J. Neurosci. Methods 76, 1–5.

Rostaing, P., Real, E., Siksou, L., Lechaire, J.-P.P., Boudier, T., Boeckers, T.M., Gertler, F., Gundelfinger, E.D., Triller, A., and Marty, S. (2006). Analysis of synaptic ultrastructure without fixative using high-pressure freezing and tomography. Eur. J. Neurosci. 24, 3463–3474.

Salin, P.A., Scanziani, M., Malenka, R.C., and Nicoll, R.A. (1996). Distinct short-term plasticity at two excitatory synapses in the hippocampus. Proc. Natl. Acad. Sci. U. S. A. 93, 13304–13309.

Shin, W., Ge, L., Arpino, G., Villarreal, S.A., Hamid, E., Liu, H., Zhao, W.-D., Wen, P.J., Chiang, H.-C., and Wu, L.-G. (2018). Visualization of Membrane Pore in Live Cells Reveals a Dynamic-Pore Theory Governing Fusion and Endocytosis. Cell 173, 934–945.e12.

Siksou, L., Varoqueaux, F., Pascual, O., Triller, A., Brose, N., and Marty, S. (2009). A common molecular basis for membrane docking and functional priming of synaptic vesicles. Eur. J. Neurosci. 30, 49–56.

De Simoni, A., Griesinger, C.B., and Edwards, F.A. (2003). Development of rat CA1 neurones in acute versus organotypic slices: role of experience in synaptic morphology and activity. J. Physiol. 550, 135–147.

Singec, I., Knoth, R., Ditter, M., Hagemeyer, C.E., Rosenbrock, H., Frotscher, M., and Volk, B. (2002). Synaptic vesicle protein synaptoporin is differently expressed by subpopulations of mouse hippocampal neurons. J. Comp. Neurol. 452, 139–153.

Smith, J.E., and Reese, T.S. (1980). Use of aldehyde fixatives to determine the rate of synaptic transmitter release. J. Exp. Biol. 89, 19–29.

Stoppini, L., Buchs, P.-A.A., and Muller, D. (1991). A simple method for organotypic cultures of nervous tissue. J. Neurosci. Methods 37, 173–182.

Studer, D., Zhao, S., Chai, X., Jonas, P., Graber, W., Nestel, S., and Frotscher, M. (2014). Capture of activity-induced ultrastructural changes at synapses by high-pressure freezing of brain tissue. Nat. Protoc. 9, 1480–1495.

Südhof, T.C.C. (2013). Neurotransmitter release: the last millisecond in the life of a synaptic vesicle. Neuron 80, 675–690.

Taschenberger, H., Woehler, A., and Neher, E. (2016). Superpriming of synaptic vesicles as a common basis for intersynapse variability and modulation of synaptic strength. Proc. Natl. Acad. Sci. 113, E4548–E4557.

Tzounopoulos, T., Janz, R., Südhof, T.C., Nicoll, R.A., and Malenka, R.C. (1998). A Role for cAMP in Long-Term Depression at Hippocampal Mossy Fiber Synapses. Neuron 21, 837–845.

Varoqueaux, F., Sigler, A., Rhee, J.-S., Brose, N., Enk, C., Reim, K., and Rosenmund, C. (2002). Total arrest of spontaneous and evoked synaptic transmission but normal synaptogenesis in the absence of Munc13-mediated vesicle priming. Proc. Natl. Acad. Sci. U. S. A. 99, 9037–9042.

Vyleta, N.P., and Jonas, P. (2014). Loose coupling between Ca2+ channels and release sensors at a plastic hippocampal synapse. Science 343, 665–670.

Vyleta, N.P., Borges-Merjane, C., and Jonas, P. (2016). Plasticity-dependent, full detonation at hippocampal mossy fiber–CA3 pyramidal neuron synapses. Elife 5.

Watanabe, S., Rost, B.R., Camacho-Pérez, M., Davis, M.W., Söhl-Kielczynski, B., Rosenmund, C., and Jorgensen, E.M. (2013). Ultrafast endocytosis at mouse hippocampal synapses. Nature 504, 242–247.

Weisskopf, M.G., Castillo, P.E., Zalutsky, R.A., and Nicoll, R.A. (1994). Mediation of hippocampal mossy fiber long-term potentiation by cyclic AMP. Science 265, 1878–1882.

Xu-Friedman, M.A., and Regehr, W.G. (2004). Structural contributions to short-term synaptic plasticity. Physiol. Rev. 84, 69–85.

